# Gradual replacement of wild bees by honeybees in flowers of the Mediterranean Basin over the last 50 years

**DOI:** 10.1101/828160

**Authors:** Carlos M. Herrera

## Abstract

Evidence for pollinator declines largely originates from mid-latitude regions in North America and Europe. Geographical heterogeneity in pollinator trends combined with geographical biases in pollinator studies, can produce distorted extrapolations and limit understanding of pollinator responses to environmental changes. In contrast to the declines experienced in some well-investigated European and North American regions, honeybees seem to have increased recently in some areas of the Mediterranean Basin. Since honeybees can impact negatively on wild bees, it was hypothesized that a biome-wide alteration in bee pollinator assemblages may be underway in the Mediterranean Basin involving a reduction in the importance of wild bees as pollinators. This hypothesis was tested using published quantitative data on bee pollinators of wild and cultivated plants from studies conducted between 1963-2017 in 13 circum-Mediterranean countries. Honeybee colonies increased exponentially and wild bees were gradually replaced by honeybees in flowers of wild and cultivated plants. Proportion of wild bees at flowers quadruplicated that of honeybees at the beginning of the period, the proportions of both groups becoming roughly similar fifty years later. The Mediterranean Basin is a world biodiversity hotspot for wild bees and wild bee-pollinated plants, and the ubiquitous rise of honeybees to dominance as pollinators could in the long run undermine the diversity of plants and wild bees in the region.

---

> *“El sur también existe”*
>
> Joan Manuel Serrat, singer and songwriter

## Introduction

The structure and dynamics of ecological communities can vary tremendously across biomes and continents. Critical elements of ecological knowledge will thus be closely tied to the particular location where it is attained, and attempts at extrapolations which are based on limited, spatially biased ecological data may produce distorted or erroneous inferences (Martin et al. 2012, Culumber et al. 2019). For instance, unawareness of geographical sampling biases has been pointed out as one possible weakness of generalizations on “pollinator decline” and “pollination crisis” (Ghazoul 2005, Archer 2014, Herrera 2019, Jamieson et al. 2019), two topics that have recently elicited considerable academic and societal interest because of the importance of animal pollination for the reproduction of many wild and crop plants (Ollerton et al. 2014, Senapathi et al. 2015, Breeze et al. 2016, Ollerton 2017). Evidence for the view of a generalized pollinator decline is strongly biased geographically, as it mostly originates from a few mid-latitude regions in Europe and North America (Rodger et al. 2004, Ghazoul 2005, Winfree et al. 2009, Archer 2014, Hung et al. 2018, Nicholson and Egan 2019). Mounting evidence indicates, however, that pollinator declines are not universal; that the sign and magnitude of temporal trends in pollinator abundance may differ among pollinator groups, continents or regions; and that taxonomic and geographical biases in pollinator studies are bound to limit a realistic understanding of the potentially diverse pollinator responses to environmental changes and the associated causal mechanisms (Aizen and Harder 2009a,b, Potts et al. 2010, vanEngelsdorp and Meixner 2010, Hofmann et al. 2018, Herrera 2019, Jamieson et al. 2019).

Even for well-studied bees, data supporting a general decline of these important pollinators tend to be geographically biased (Archer et al. 2014, Ollerton 2017, Hung et al. 2018). For example, in thoroughly studied North America and mid-western Europe the number of honeybee (*Apis mellifera*) colonies has experienced severe declines, but the trend is apparently reversed in the less investigated areas of southern Europe, where honeybee colonies seem to have been steadily increasing over large territories in the last decades (Aizen and Harder 2009a: Fig. S1, Potts et al. 2010, vanEngelsdorp and Meixner 2010, Moritz and Erler 2016). Honeybees have been repeatedly shown to have negative impacts on wild bee populations in both natural and anthropogenic scenarios (Goulson and Sparrow 2009, Shavit et al. 2009, Lindström et al. 2016, Torné-Noguera et al. 2016, Magrach et al. 2017, Ropars et al. 2019, Valido et al. 2019). I thus formulated the hypothesis that, if the abundance of managed honeybees has been actually increasing in the Mediterranean Basin over the last decades, then a profound biome-wide alteration in the proportional composition of bee pollinator assemblages could be currently underway there, involving a gradual replacement of wild bees by honeybees in flowers. This paper verifies this hypothesis using data from a large sample of published investigations on the bee pollinators of wild and cultivated plants, conducted during the last 50 years throughout the Mediterranean Basin. Results of this study stress the importance of broadening the geographical scope of current investigations on pollinator trends, while at the same time issue a warning on the perils of uncritically importing to Mediterranean ecosystems honeybee conservation actions specifically designed for the contrasting situations that prevail in temperate-climate European or North American countries.

## Material and methods

### The data

The literature on floral biology, pollination ecology, plant-pollinator interactions and crop pollination was searched for field studies conducted during 1960-2019 in the Mediterranean Basin and providing quantitative data on the relative abundance of honeybees and wild bees at flowers of insect-pollinated plants, either wild-growing or cultivated. Preliminary searches had shown that studies conducted before 1960 quite rarely reported quantitative data on bee abundance at flowers. The literature screening used searches in Web of Science, Google Scholar and my personal database of plant-pollinator studies. To improve the chances of obtaining a representative, geographically comprehensive coverage of all regions surrounding the Mediterranean Sea (i.e., African, Asian and European shores), literature searches were conducted using terms in English, French, Italian, Portuguese and Spanish. For inclusion in this study I considered exclusively field investigations where (1) quantitative data were provided on numbers or relative proportions of wild bee and honeybee individuals recorded at flowering plants or flowering patches of single plant species, obtained using direct visual counts or standardized collections. Investigations at the plant community level or providing semiquantitative or subjective abundance scores of bee abundance were thus excluded; and (2) the year(s) on which bee abundance data had been originally collected in the field was unambiguously stated. In a few publications where information from two or more study years had been pooled into a single estimate of wild bee and honeybee abundances, but the data were otherwise suitable, the average year was used. A total of 336 estimates of wild bee and honeybee abundance at the flowers of 200 plant species were gathered from 136 different literature sources. Original figures of bee abundance at flowers were transformed to proportions of wild bee (*p*_wb_) and honeybee (*p*_hb_ = 1 - *p*_wb_) individuals relative to individuals of all bees combined. Each data record corresponded to a unique combination of plant species x sampling year x sampling location. The data had been collected in the field between 1963–2017 in 13 different countries surrounding the Mediterranean Sea (Fig. 1). Information on plant type (wild-growing *vs.* cultivated) and taxonomic affiliation (plant family) was also incorporated into the data set.

**Fig. 1.**
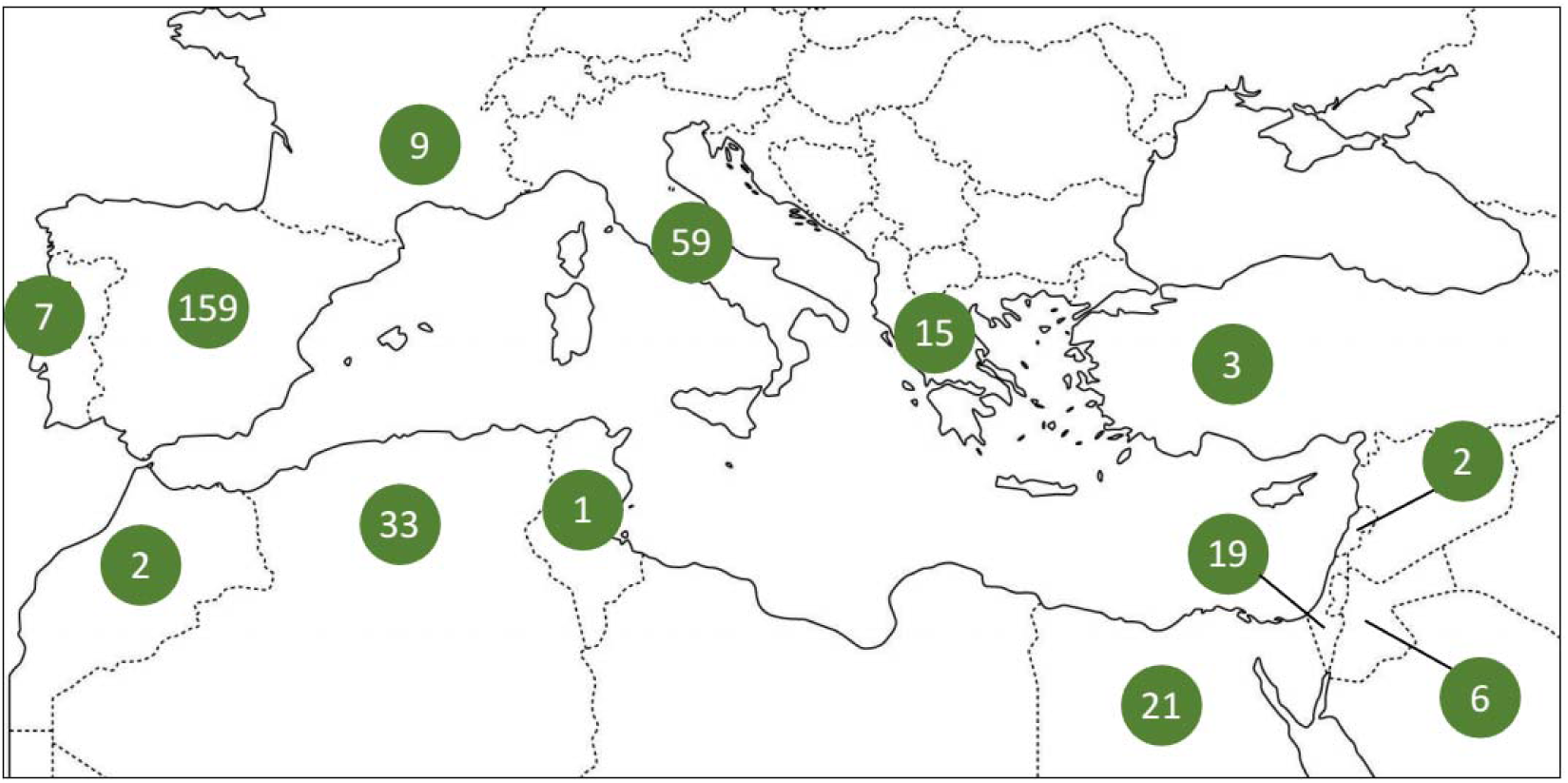
Distribution among 13 circum-Mediterranean countries of the *N* = 336 published estimates of wild bee and honeybee abundance in flowers of cultivated and wild-growing plants for the period 1963-2017 considered in this study (Table S1, electronic supplementary material).

The complete data set including literature sources is presented in Table S1, electronic supplementary material. Most data originated from Spain, Italy, Algeria and Egypt (159, 59, 33 and 21 records, respectively; Fig. 1). The median of the distribution of study years was 1996 (interquartile range = 1986-2008). There were 106 and 230 records for cultivated and wild-growing plants, respectively. A total of 54 plant families were represented in the sample, with most species belonging to Fabaceae, Lamiaceae, Asteraceae, Rosaceae and Cistaceae; 51, 34, 32, 30 and 25 records, respectively).

Trends in honeybee abundance in the Mediterranean Basin over the period considered in this study were assessed using information gathered from the Food and Agriculture Organization (FAO) of the United Nations global database (FAOSTAT;http://www.fao.org/faostat). This data source has been used previously in historical reviews of honeybee abundance (Aizen and Harder 2009a,b, vanEngelsdorp and Meixner 2010, Moritz and Erler 2016). Number of honeybee colonies per country and year for the period 1963-2017 was obtained from FAOSTAT (accessed 25 September 2019) for each of the 13 Mediterranean countries with estimates of wild bee and honeybee relative abundances in my data set (Fig. 1). Comparable abundance figures were obtained by dividing the number of honeybee colonies by the land surface of the country (obtained also from FAOSTAT), which provided estimates of honeybee colonies/km^2^ per country and year. Data on honey production per country and year were also obtained from FAOSTAT to check the reliability of colony numbers as a suitable proxy for honeybee abundance (Aizen and Harder 2009b, Moritz and Erler 2016).

### Statistical analyses

For the purpose of statistical analyses, the log-odds that one randomly chosen bee found at flowers was a wild bee rather than a honeybee was estimated for each data record using the logit transformation, logit(*p*_wb_) = log(*p*_wb_/*p*_hb_). Since the logit function is undefined for *p* = 0 or 1, proportions were remaped to the interval (0.05, 0.95) prior to the transformation.

The null hypothesis that the relative proportions of wild bees and honeybees at flowers were unrelated to year of data collection was tested by fitting a linear mixed effect model. Logit(*p*_wb_) was the response variable, and data collection year (treated as a continuous numerical variable), plant type (two-level factor, wild-growing *vs.* cultivated) and their interaction were included as fixed effects. Country of origin, plant family and plant species were included as random effects to statistically control for, on one side, the effects of likely taxonomic and geographical correlations in the data and, on the other, the unbalanced distribution of data across countries and plant taxonomic groups. The existence of a long-term trend in honeybee abundance in the Mediterranean Basin as a whole was tested by fitting a linear mixed model to the FAOSTAT colony density data (log-transformed). Year (as a numerical variable) was the single fixed effect, and country was included in the model as a random effect to account for the correlated data of the same country. Linear mixed models allow drawing conclusions on fixed effects with reference to a broad inference space whose scope transcends the specific samples studied (McLean et al. 1991, Bolker 2015). In the present instance, the universe of all countries and plant species in the Mediterranean Basin that could have been sampled for this study represents the broad inference space (Schabenberger and Pierce 2001). Conclusions on long-term trends in honeybee abundance and logit(*p*_wb_), including predicted marginal effects, will thus refer to such broad inference space.

All statistical analyses were carried out using the R environment (R Core Team 2018). Linear mixed models were fitted with the lmer function in the lme4 package (Bates et al. 2015). Statistical significance of fixed effects was assessed using analysis of deviance-based, Type II Wald Chi-square tests using the Anova function in the car package (Fox and Weisberg 2011). The function ggpredict from the ggeffects package (Lüdecke 2018) was used to compute marginal effects of year on logit(*p*_wb_) separately for wild-growing and cultivated plants.

## Results

Estimated density of managed honeybee colonies tended to increase steadily over the 1963-2017 period in most Mediterranean countries considered in this paper (Fig. 2). The linear mixed model fitted to colony density data (log-transformed), with year as fixed effect and country as random effect, revealed a highly significant, positive linear effect of year on colony density (Chi-squared = 412.9, *P* < 10^−16^). The estimated linear trend for the whole Mediterranean Basin obtained from this model is depicted in Fig. 2. Linearity of the estimated relationship on the logarithmic scale reveals an exponential increase in the density of honeybee colonies in the region as a whole over the period considered. There was a close linear relationship across years between mean honey production and mean number of honeybee colonies per country and year (Fig. S1, electronic supplementary material), which supports the reliability of FAOSTAT colony number data as a proxy for honeybee abundance.

**Fig. 2.**
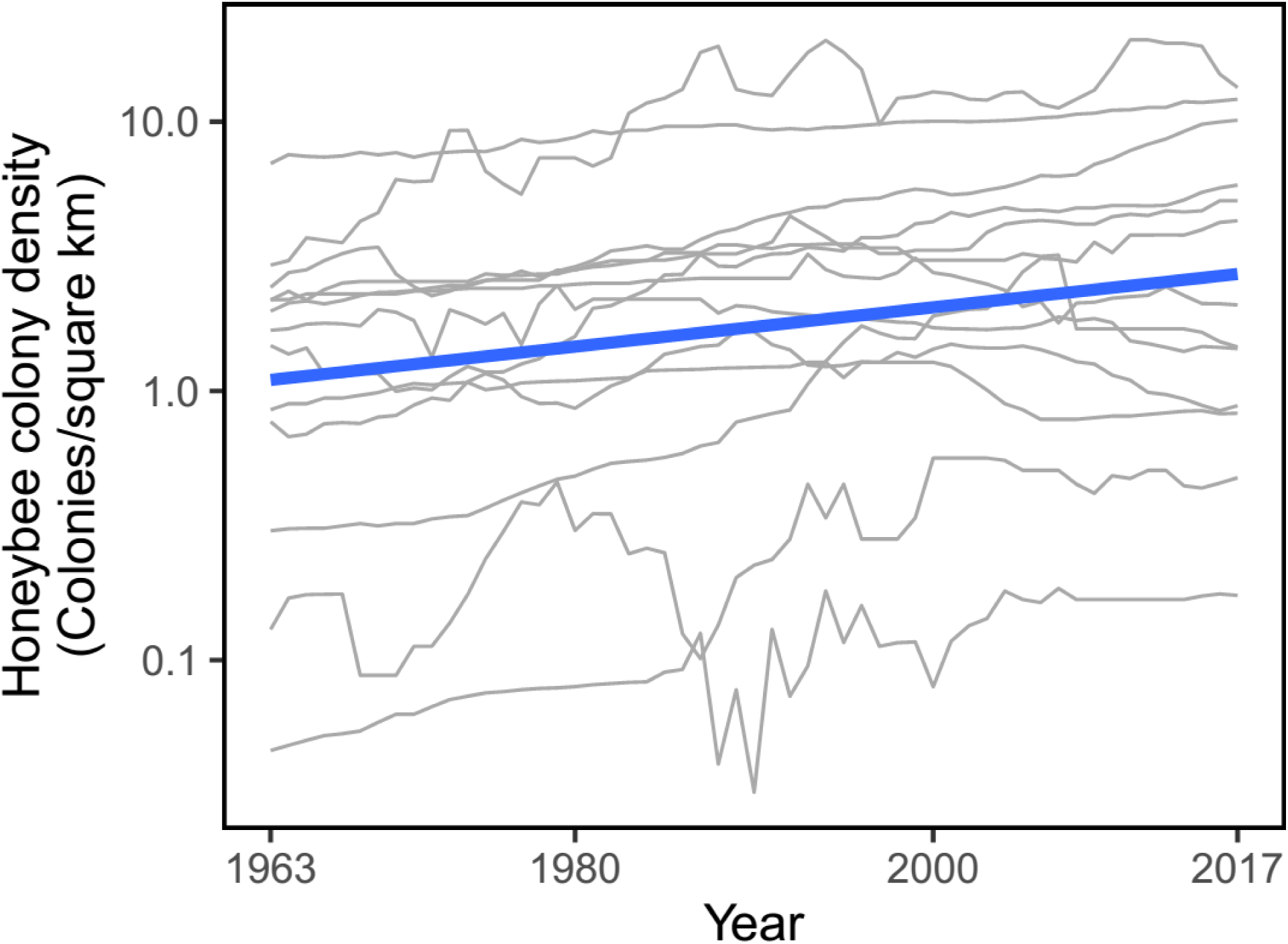
Variation over 1963-2017 in density of honeybee colonies in the 13 circum-Mediterranean countries considered in this study (gray lines), and overall relationship for the Mediterranean Basin as a whole (blue line), as estimated from parameters of a linear mixed model fitted to the data with country as a random effect. Note the logarithmic scale on vertical axis.

For all years, countries and plant species combined, the logarithm of the ratio between proportions of wild bees and honeybees at flowers [logit(*p*_wb_)] encompassed the whole range of possible values, and there was extensive overlap between cultivated and wild-growing plants (Fig. 3). Wild bees tended to be proportionally more abundant in flowers of wild-growing plants (mean logit(*p*_wb_) ± SE = 0.655 ± 0.120, *N* = 230) than in flowers of cultivated ones (–0.242 ± 0.167, *N* = 106), the difference being statistically significant (Chi-squared = 18.96, *P* = 0.000013, Kruskal-Wallis rank sum test). For all the data combined (“naïve” least-squares regression fitted to the data; Fig. 4A), there existed a statistically significant, negative relationship between logit(*p*_wb_) and year of study (*r*_s_ = - 0.139, *N* = 336, *P* = 0.011, Spearman rank correlation), thus suggesting a declining trend in the importance of wild bees at flowers relative to honeybees over the period considered (Fig. 4A). The reality of this trend was corroborated and strenghtened after statistically accounting for correlations underlying the data and the unbalanced distribution across plant types, countries, plant families and plant species.

**Fig. 3.**
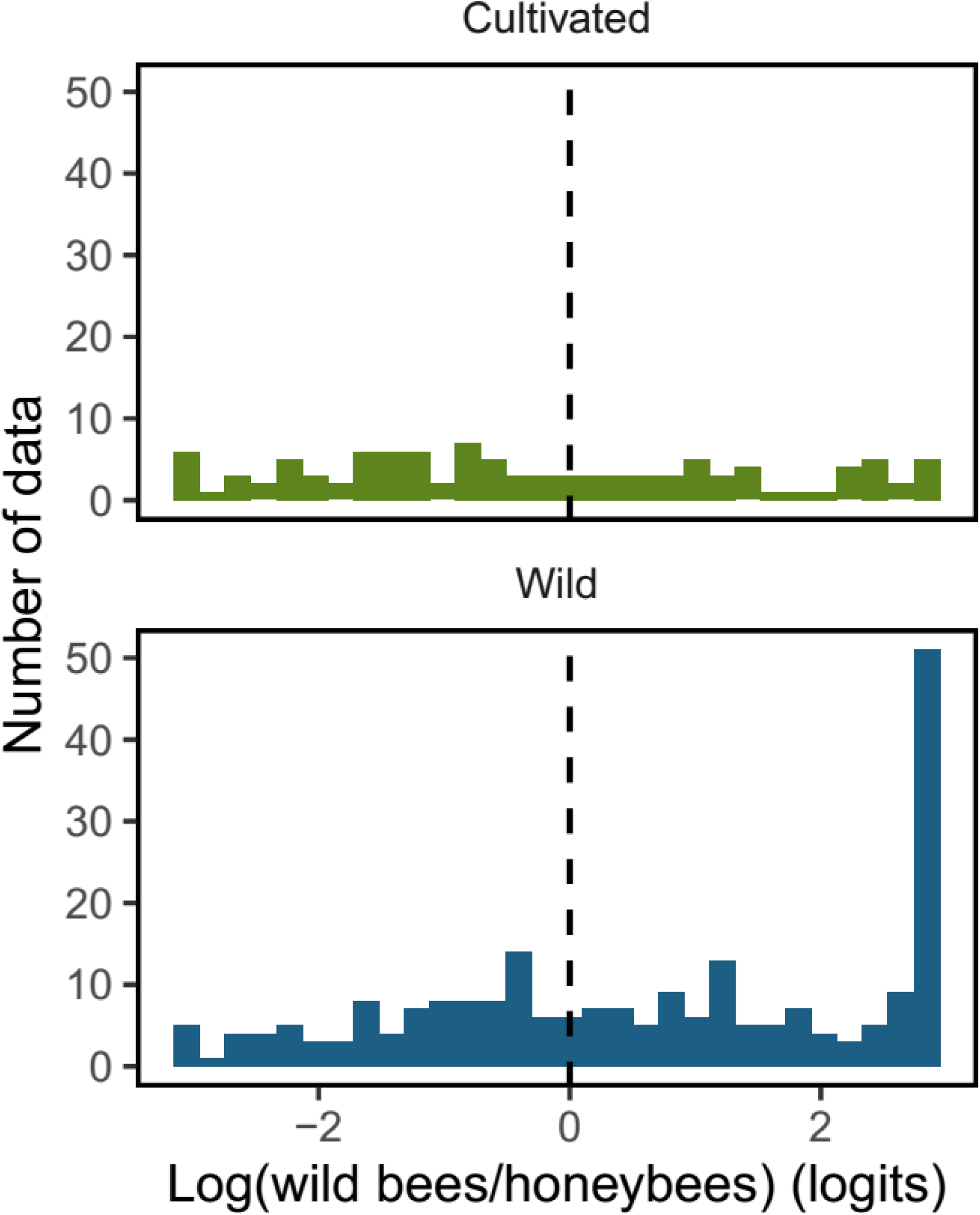
Frequency distributions of logit(*p*_wb_), the logarithm of the ratio between proportions of wild bees and honeybees at flowers, in the *N* = 336 unique combinations of plant species x sampling year x sampling location considered in this study (*N* = 106 and 230 records for cultivated and wild plants, respectively). Bars to the left and right of the vertical dashed line [logit(*p*_wb_) = 0] correspond to situations of numerical dominance at flowers of honeybees and wild bees, respectively.

**Fig. 4.**
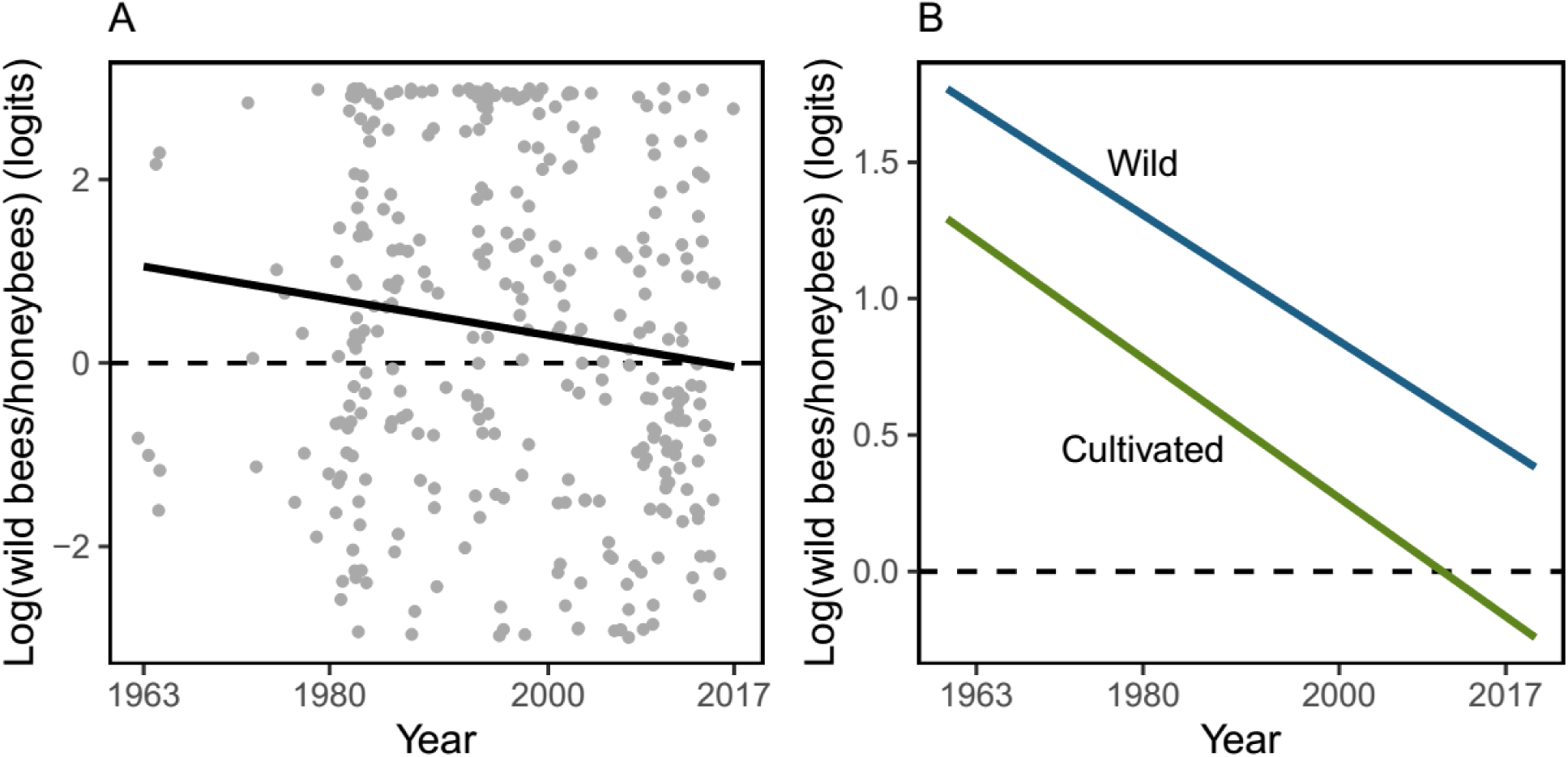
A. Relationship between logit(*p*_wb_), the logarithm of the ratio between proportions of wild bees and honeybees at flowers, and year of study. Each dot corresponds to a unique combinations of plant species x sampling year x sampling location (*N* = 336). The black line is the “naïve” least-squares regression fitted to the data, all countries, plant species and plant types (cultivated and wild-growing) combined. B. Mean estimated marginal effects of year on logit(*p*_wb_) for cultivated and wild-growing plants, as predicted from the linear mixed model with country, plant family, and plant species as random effects (Table 1).

Results of the linear mixed model testing for the effect of year of study on logit(*p*_wb_) are summarized in Table 1. After statistically accounting for plant type (wild-growing *vs.* cultivated), country, plant family and plant species, there was a highly significant negative effect of study year on logit(*p*_wb_). The effect was similar for wild-growing and cultivated species, as denoted by the statistical nonsignificance of the year x plant type interaction. The effect of plant type on logit(*p*_wb_) was only marginally significant after statistically accounting for the rest of effects in the model (Table 1). Mean predicted marginal effects of year on logit(*p*_wb_), computed separately for wild-growing and cultivated plants, illustrate a linear decline in logit(*p*_wb_) over the study period (Fig. 4B). The data-predicted proportion of wild bees at flowers for 1963 roughly quadruplicated that of honeybees, while the predicted proportions of both groups for 2017 were roughly similar. This long-term replacement of wild bees by honeybees at flowers occurred at similar rates in wild and cultivated plants, as shown by the parallel predicted marginal effects (Fig. 4B).

**Table 1.** Summary of results of the linear mixed model testing for the significance of supra-annual variation in logit(*p*_wb_), the log of the quotient between proportions of wild bees and honeybees, in flowers of wild-growing and cultivated plants of the Mediterranean Basin.

|  | Standardized<br>parameter estimate<br>(standard error) | Chi-squared | *P* value | Variance<br>(95% confidence<br>interval) |
| --- | --- | --- | --- | --- |
| Fixed effects |  |  |  |  |
| Year (Y) | -0.314 (0.137) | 10.94 | 0.00094 |  |
| Plant type (PT) | 0.566 (0.306) | 3.45 | 0.063 |  |
| Y x PT | 0.030 (0.184) | 0.027 | 0.87 |  |
| Random effects |  |  |  |  |
| Country |  |  |  | 0.357 (0.040-1.254) |
| Plant family |  |  |  | 0.389 (0.091-0.983) |
| Plant species |  |  |  | 1.399 (0.896-1.997) |

## Discussion

Previous studies that have examined long-term trends in honeybee colony numbers from a wide geographical perspective have consistently shown that (1) there is not any hint of honeybees declining at a planetary scale, but rather considerable evidence that the total number of colonies is increasing globally and in almost every continent; (2) well-documented instances of honeybee decline are few and fairly restricted geographically, being mostly circumscribed to parts of Europe and North America; and (3) in the thoroughly-investigated European continent, honeybee declines have occurred in mid-latitude and northern countries, while increases predominate in the south (Aizen and Harder 2009a, Potts et al. 2010, vanEngelsdorp and Meixner 2010, Moritz and Erler 2016). As an example, Fig. 5 depicts the opposite trajectories of honeybee colony density over the last half century in two countries representative from mid-western Europe and the Mediterranean Basin (see also vanEngelsdorp and Meixnar: Fig. 2). The analyses presented in this study show that honeybee colonies have increased exponentially over the last 50 years in the Mediterranean Basin, comprising areas of southern Europe, the Middle East and Northern Africa. The latter two regions are prominent examples of ecologically understudied areas (Martin et al. 2012) and, as far as I know, have been never considered in quantitative analyses of bee population trends. The empirical evidence available, therefore, supports the view that to the extent that extrapolations on “pollinator decline” or “pollination crisis” were at some time inspired by the decline of honeybees in a few regions (see, e.g., Ghazoul 2005, Potts et al. 2010, Ollerton 2017, for reviews), such generalizations represent prime examples of distorted ecological knowledge arising from geographically biased data (Ghazoul 2005, Martin et al. 2012, Archer et al. 2014, Culumber et al. 2019).

**Fig. 5.**
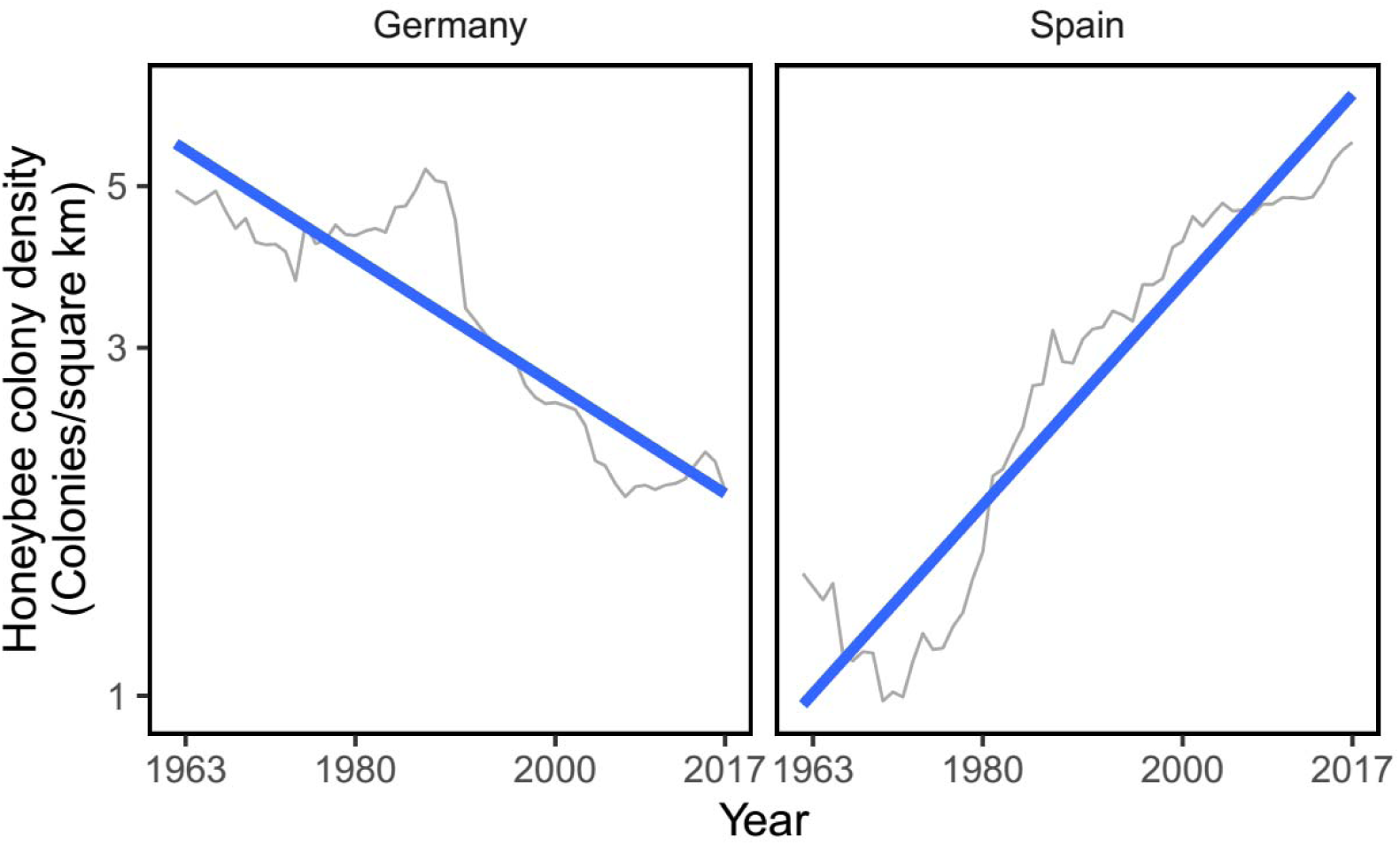
Variation over 1963-2017 in density of honeybee colonies in Germany an Spain (gray lines), based on FAOSTAT data (see text). These two countries were chosen as representatives, respectively, of thoroughly-studied, mid-western, temperate-climate Europe, and insufficiently-studied, southern, Mediterranean-climate Europe. Blue lines represent least-squares fitted linear regressions.

Correlative and experimental evidence alike has shown that at the local and regional scales honeybees can have strong negative impacts on wild bee populations in both natural and anthropogenic scenarios (Shavit et al. 2009, Lindström et al. 2016, Torné-Noguera et al. 2016, Magrach et al. 2017, Ropars et al. 2019, Valido et al. 2019), and that the absence of honeybees in well-preserved natural areas is associated with increasing wild bee populations (Herrera 2019). Much of the direct or circumstantial evidence on the harmful effects of honeybees on wild bees originated in the Mediterranean Basin, which motivated the hypothesis formulated in this paper of a possible replacement of wild bees by honeybees in the Mediterranean in parallel to increasing honeybee abundance. This hypothesis has been tested using literature data from highly heterogeneous sources, and originally collected using an enormous variety of field procedures. The data were also imbalanced with regard to observation year, country of origin or plant taxonomic affiliation, all of which combined to produce a “messy” dataset. Despite these limitations of the data, the prediction of a gradual long-term replacement of wild bees by honeybees in flowers of the Mediterranean Basin was verified. This conclusion persisted regardless of whether the hypothesis was tested “naïvely” (i.e., simple correlation on all data pooled) or by fitting a linear mixed model where major sources of data “messiness” were appropriately handled by treating them as random effects. Estimated marginal effects predicted from the mixed model revealed that, on average, the proportion of wild bees at Mediterranean flowers roughly quadruplicated that of honeybees at the beginning of the period considered (logit(*p*_wb_) ∼ 1.5) while fifty years later the proportions of the two groups had become roughly similar (logit(*p*_wb_) ∼ 0).

On average, model-predicted importance of wild bees relative to honeybees was slightly lower in flowers of cultivated plants throughout the period considered, a finding that seems logically related to the traditional practice of placing honeybee colonies in the vicinity of orchards or cultivated land to ensure crop pollination. More difficult to interpret is the close similarity between wild and cultivated plants in average replacement rate of wild bees by honeybees in flowers, denoted by parallel slopes of mean predicted marginal effects of year on logit(*p*_wb_) and statistical nonsignificance of the year x plant type interaction effect. A tentative interpretation of this finding is that the causal mechanism behind trends in the proportional composition of bees at flowers was one and the same for cultivated and wild plants, or in other words, that increasing honeybee colony density induced similar proportional reductions of wild bees in flowers from anthropogenous and natural habitats. Irrespective of the causal mechanism accounting for it, however, parallel trends in the proportional decline of wild bees relative to honeybees in wild and cultivated plants corroborate in a broader geographical context previous findings at a regional scale showing that natural Mediterranean habitats are not exempt from the negative impact of increasing honeybee densities in anthropogenous habitats nearby (Magrach et al. 2017).

Results of this study are important because the Mediterranean Basin is a world biodiversity hotspot for both wild bees and wild bee-pollinated plants (Petanidou and Vokou 1993, Dafni and O’Toole 1994, Michener 2000, Petanidou and Lamborn 2005, Harrison and Noss 2017). Predicting the global consequences for the Mediterranean flora of the proportional decline of wild bees as floral visitors documented in this paper will require extensive data, e.g., on the pollinating effectiveness of different groups of bees on different plants. Nevertheless, studies conducted so far on the effectiveness of honeybees and wild bees as pollinators of cultivated and wild species in the Mediterranean Basin have shown that wild bees generally are better pollinators than honeybees (Herrera 1987, Obeso 1992, Bosch and Blas 1994, Vicens and Bosch 2000, Potts et al. 2001, Monzón et al. 2004). If these limited findings are corroborated in the future by more extensive investigations, then the gradual replacement of wild bees by honeybees currently underway in Mediterranean flowers could translate into impaired fruit and seed production and, in the case of pollen-limited wild plants, reduced population recruitment.

It does not seem implausible to suggest that, because of its colossal magnitude and spatial extent, the exponential flood of honeybee colonies that is silently taking over the Mediterranean Basin can pose serious threats to two hallmarks of the Mediterranean biome, namely the extraordinary diversities of wild bees and wild bee-pollinated plants (Blondel et al. 2010). The Mediterranean Basin is home to ∼3300 wild bee species, or ∼87% of those occurring in the whole Western Palaearctic region (data from Discover Life, https://www.discoverlife.org/, accessed 1 November 2019; and Kuhlmann 2019). Large as that percentage may seem, it is likely an underestimate given the imperfect knowledge of the rich bee faunas of Mediterranean Africa and Asia. From a conservation perspective, the technical, political and administrative actions launched for promoting apiculture or enhancing honeybee populations in those European regions where the species is declining (European Parliament 2008, de la Rúa et al. 2009, Cayuela et al. 2011) should not be transferred uncritically to the Mediterranean Basin. In the Mediterranean countries, such actions would not only be aiming at the wrong conservation target but, much worse, could be inadvertently threatening the unique regional diversity of wild bees, wild bee-pollinated plants and their mutualistic relationships.

## Acknowledgements

This study was prompted by the troubling discrepancy between allusions to the honeybees’ impending demise so often found in popular media, and my strong subjective impression in the field that managed honeybees are displacing wild bees from flowers in the Iberian Peninsula. Assistance from the Red de Bibliotecas y Archivos del CSIC was essential for procuring old publications from rather oscure pre-Internet resources. I am grateful to Oscar Aguado, Angel Guardiola, Fernando Jubete and Alejandro Martínez Abraín for stimulating discussion, and Conchita Alonso and Mónica Medrano for useful suggestions on the manuscript. The research reported in this paper received no specific grant from any funding agency.

