## Supplementary material for "Gradual replacement of wild bees by honeybees in flowers of the Mediterranean Basin over the last 50 years": Fig. S1

Figure S1. Relationship across years over the period 1963-2017 between mean per-country honey production and number of honeybee colonies for the set of *N* = 13 circum-Mediterranean countries considered. Each dot represents the average values of honey production and honeybee colonies in a given year, and the blue line is the least-squares fitted regression. Data from the Food and Agriculture Organization (FAO) of the United Nations global database (FAOSTAT; http://www.fao.org/faostat).

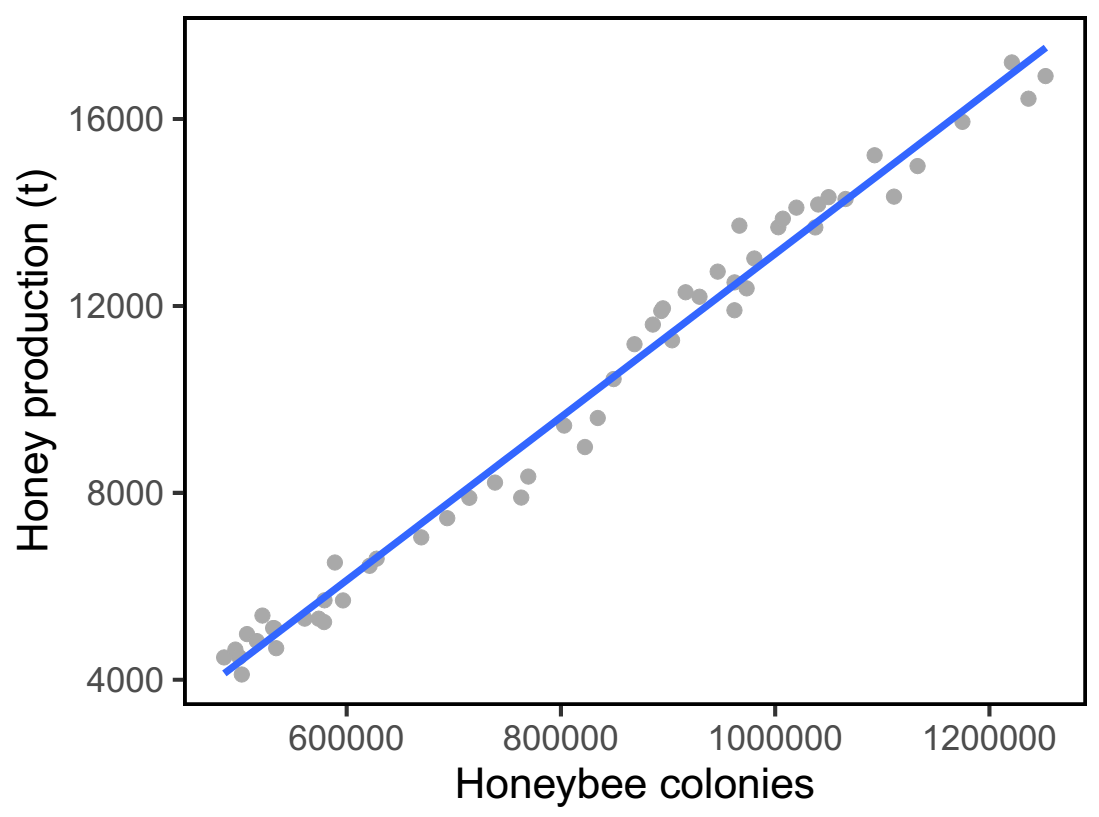
