## Supplemental Data 1 for "Gradual replacement of wild bees by honeybees in flowers of the Mediterranean Basin over the last 50 years"

| Table S1. Raw data used in the study and literature references. | | | | | | |
| --- | --- | --- | --- | --- | --- | --- |
| Species | Family | Plant type | Country | Year of study | Wild bees relative to total bees (%) | References * |
| *Actinidia deliciosa* | Actinidiaceae | Cultivated | Spain | 2013 | 42.66 | Miñarro and Twizell (2015) |
| *Agastache foeniculum* | Labiatae | Cultivated | Italy | 1992 | 8.08 | Quaranta and Ricciardelli d'Albore (1996) |
| *Agastache foeniculum* | Labiatae | Cultivated | Italy | 1993 | 15.16 | Quaranta and Ricciardelli d'Albore (1996) |
| *Allium porrum* | Alliaceae | Cultivated | Italy | 2015 | 28.40 | Fijen and Kleijn (2017) |
| *Anacyclus clavatus* | Asteraceae | Wild | Algeria | 1994 | 100.00 | Louadi and Doumandji (1998) |
| *Andryala integrifolia* | Asteraceae | Wild | Spain | 1986 | 87.24 | Bosch et al. (1997) |
| *Angelica archangelica* | Apiaceae | Wild | Italy | 1984 | 67.76 | Ricciardelli d'Albore (1986) |
| *Anthyllis barba-jovis* | Fabaceae | Wild | Italy | 2015 | 14.20 | Benelli et al. (2017) |
| *Anthyllis vulneraria* | Fabaceae | Wild | Spain | 1993 | 100.00 | Navarro (2000) |
| *Anthyllis vulneraria* | Fabaceae | Wild | Spain | 1994 | 100.00 | Navarro (2000) |
| *Anthyllis vulneraria* | Fabaceae | Wild | Spain | 1996 | 100.00 | Navarro (2000) |
| *Antirrhinum microphyllum* | Plantaginaceae | Wild | Spain | 1997 | 100.00 | Torres et al. (2001) |
| *Apium graveolens* | Apiaceae | Cultivated | Italy | 1984 | 99.04 | Ricciardelli d'Albore (1986) |
| *Armeria velutina* | Plumbaginaceae | Wild | Spain | 1982.5 | 100.00 | J. Herrera (1985, 1988) |
| *Asparagus aphyllus* | Asparagaceae | Wild | Spain | 1982.5 | 100.00 | J. Herrera (1985, 1988) |
| *Asphodelus albus* | Asphodelaceae | Wild | Spain | 1988 | 30.40 | Obeso (1992) |
| *Asphodelus albus* | Asphodelaceae | Wild | Spain | 1989 | 76.20 | Obeso (1992) |
| *Asphodelus albus* | Asphodelaceae | Wild | Spain | 1990 | 97.70 | Obeso (1992) |
| *Astragalus glycyphyllos* | Fabaceae | Cultivated | Italy | 1983 | 14.08 | Ricciardelli d'Albore (1983) |
| *Astragalus tragacantha* | Fabaceae | Wild | France | 2017 | 98.82 | Schurr et al. (2019) |
| *Atropa bella-donna* | Solanaceae | Wild | Italy | 1983 | 10.47 | Ricciardelli d'Albore (1984) |
| *Beta vulgaris* | Amaranthaceae | Cultivated | Algeria | 2004 | 97.30 | Benachour (2008) |
| *Borago officinalis* | Boraginaceae | Wild | Algeria | 1994 | 91.30 | Louadi and Doumandji (1998) |
| *Borago officinalis* | Boraginaceae | Wild | Spain | 2014 | 12.83 | Pérez-Marcos et al. (2017) |
| *Brassica fruticulosa* | Brassicaceae | Wild | Spain | 1986 | 79.70 | Bosch et al. (1997) |
| *Brassica napus* | Brassicaceae | Cultivated | Egypt | 2014 | 81.40 | Kamel et al. (2015) |
| *Brassica napus* | Brassicaceae | Cultivated | Egypt | 2015 | 73.00 | Kamel et al. (2015) |
| *Brassica oleracea* | Brassicaceae | Cultivated | Egypt | 1981 | 2.10 | Hussein and Abdel-Aal (1982) |
| *Brassica oleraceae* | Brassicaceae | Wild | Spain | 2014 | 73.90 | Pérez-Marcos et al. (2017) |
| *Brassica rapa* | Brassicaceae | Cultivated | Algeria | 2003 | 14.23 | Benachour (2008) |
| *Brassica sinapis* | Brassicaceae | Cultivated | Egypt | 1981 | 19.20 | Hussein and Abdel-Aal (1982) |
| *Buxus balearica* | Buxaceae | Wild | Spain | 2002 | 2.11 | Lázaro and Traveset (2005) |
| *Buxus balearica* | Buxaceae | Wild | Spain | 2002 | 13.74 | Lázaro and Traveset (2005) |
| *Buxus balearica* | Buxaceae | Wild | Spain | 2003 | 4.02 | Lázaro and Traveset (2005) |
| *Buxus balearica* | Buxaceae | Wild | Spain | 2003 | 14.46 | Lázaro and Traveset (2005) |
| *Buxus balearica* | Buxaceae | Wild | Spain | 2003 | 40.86 | Lázaro and Traveset (2005) |
| *Buxus balearica* | Buxaceae | Wild | Spain | 2003 | 60.00 | Lázaro and Traveset (2005) |
| *Calendula arvensis* | Asteraceae | Wild | Algeria | 1994 | 100.00 | Louadi and Doumandji (1998) |
| *Calendula suffruticosa* | Asteraceae | Wild | Algeria | 1994 | 99.29 | Louadi and Doumandji (1998) |
| *Calluna vulgaris* | Ericaceae | Wild | Spain | 1982.5 | 82.61 | J. Herrera (1985, 1988) |
| *Calluna vulgaris* | Ericaceae | Wild | France | 2013 | 17.39 | Descamps et al. (2015) |
| *Campanula lingulata* | Campanulaceae | Wild | Greece | 1993 | 100.00 | Blionis and Vokou (2001) |
| *Campanula lingulata* | Campanulaceae | Wild | Greece | 1994 | 100.00 | Blionis and Vokou (2001) |
| *Campanula lingulata* | Campanulaceae | Wild | Greece | 1994 | 100.00 | Blionis and Vokou (2001) |
| *Campanula oreadum* | Campanulaceae | Wild | Greece | 1994 | 80.04 | Blionis and Vokou (2001) |
| *Campanula persicifolia* | Campanulaceae | Wild | Greece | 1994 | 100.00 | Blionis and Vokou (2001) |
| *Campanula rotundifolia* | Campanulaceae | Wild | Greece | 1994 | 34.70 | Blionis and Vokou (2001) |
| *Campanula spatulata* | Campanulaceae | Wild | Greece | 1994 | 100.00 | Blionis and Vokou (2001) |
| *Campanula versicolor* | Campanulaceae | Wild | Greece | 1994 | 39.08 | Blionis and Vokou (2001) |
| *Campanula versicolor* | Campanulaceae | Wild | Greece | 1994 | 78.96 | Blionis and Vokou (2001) |
| *Capparis ovata* | Capparaceae | Wild | Israel | 1982 | 38.03 | Dafni et al. (1987) |
| *Capparis spinosa* | Capparaceae | Wild | Israel | 1982 | 30.30 | Dafni et al. (1987) |
| *Carthamus tinctorius* | Asteraceae | Cultivated | Italy | 1981 | 18.60 | Frediani and Pinzauti (1985) |
| *Carum carvi* | Apiaceae | Cultivated | Italy | 1984 | 98.29 | Ricciardelli d'Albore (1986) |
| *Centaurea aspera* | Asteraceae | Wild | Spain | 1986 | 41.80 | Bosch et al. (1997) |
| *Centaurea cyanus* | Asteraceae | Wild | France | 2001 | 80.61 | Vanderbergh (2002) |
| *Centaurea jacea* | Asteraceae | Wild | France | 2001 | 60.28 | Vanderbergh (2002) |
| *Centaurea scabiosa* | Asteraceae | Wild | France | 2001 | 71.70 | Vanderbergh (2002) |
| *Centaurea solstitialis* | Asteraceae | Wild | Greece | 2008 | 100.00 | Barthell et al. (2012) |
| *Cephalanthera longifolia* | Orchidaceae | Wild | Israel | 1978.5 | 100.00 | Dafni and Ivri (1981) |
| *Cephalaria transsylvanica* | Dipsacaceae | Wild | Italy | 2013 | 74.80 | Benelli et al. (2014) |
| *Chrysanthemum coronarium* | Asteraceae | Wild | Spain | 2014 | 93.67 | Pérez-Marcos et al. (2017) |
| *Chrysanthemum paludosum* | Asteraceae | Wild | Algeria | 1994 | 98.53 | Louadi and Doumandji (1998) |
| *Cistus albidus* | Cistaceae | Wild | Spain | 1985 | 72.60 | Bosch (1992) |
| *Cistus creticus* | Cistaceae | Wild | Greece | 1998 | 20.00 | Manetas and Petropoulou (2000) |
| *Cistus creticus* | Cistaceae | Wild | Greece | 2013.5 | 87.02 | Goras et al. (2016) |
| *Cistus crispus* | Cistaceae | Wild | Spain | 2011 | 24.30 | Magrach et al. (2017) |
| *Cistus crispus* | Cistaceae | Wild | Spain | 2012 | 38.30 | González-Varo et al. (2016) |
| *Cistus crispus* | Cistaceae | Wild | Spain | 2012 | 36.47 | Magrach et al. (2017) |
| *Cistus ladanifer* | Cistaceae | Wild | Spain | 2011 | 18.87 | Magrach et al. (2017) |
| *Cistus ladanifer* | Cistaceae | Wild | Spain | 2012 | 33.02 | Magrach et al. (2017) |
| *Cistus libanotis* | Cistaceae | Wild | Spain | 1982.5 | 93.08 | J. Herrera (1985, 1988) |
| *Cistus libanotis* | Cistaceae | Wild | Spain | 1996 | 100.00 | Talavera et al. (2001) |
| *Cistus monspeliensis* | Cistaceae | Wild | Spain | 1985 | 87.50 | Bosch (1992) |
| *Cistus monspeliensis* | Cistaceae | Wild | Spain | 2011 | 28.30 | Magrach et al. (2017) |
| *Cistus monspeliensis* | Cistaceae | Wild | Spain | 2012 | 39.83 | Magrach et al. (2017) |
| *Cistus salviifolius* | Cistaceae | Wild | Spain | 1982.5 | 34.18 | J. Herrera (1985, 1988) |
| *Cistus salviifolius* | Cistaceae | Wild | Spain | 1985 | 97.40 | Bosch (1992) |
| *Cistus salviifolius* | Cistaceae | Wild | Spain | 2011 | 34.42 | Magrach et al. (2017) |
| *Cistus salviifolius* | Cistaceae | Wild | Spain | 2012 | 23.31 | Magrach et al. (2017) |
| *Citrullus lanatus* | Cucurbitaceae | Cultivated | Egypt | 2006 | 8.60 | Taha and Bayoumi (2009) |
| *Colchicum stevenii* | Colchicaceae | Wild | Israel | 1988 | 83.33 | Dafni (1996) |
| *Convolvulus arvensis* | Convolvulaceae | Wild | Algeria | 1994 | 89.47 | Louadi and Doumandji (1998) |
| *Convolvulus tricolor* | Convolvulaceae | Wild | Algeria | 1994 | 77.78 | Louadi and Doumandji (1998) |
| *Coriandrum sativum* | Apiaceae | Cultivated | Egypt | 1981 | 32.30 | Hussein and Abdel-Aal (1982) |
| *Coriandrum sativum* | Apiaceae | Cultivated | Jordan | 2007 | 0.00 | Abu-Hammour (2008) |
| *Coriandrum sativum* | Apiaceae | Cultivated | Algeria | 2009 | 82.40 | Bendifallah et al. (2013) |
| *Coriandrum sativum* | Apiaceae | Wild | Spain | 2014 | 22.27 | Pérez-Marcos et al. (2017) |
| *Cornus sanguinea* | Cornaceae | Wild | Spain | 1993 | 56.60 | Guitián et al. (1996) |
| *Crataegus monogyna* | Rosaceae | Wild | Spain | 1988.5 | 19.13 | Guitián and Fuentes (1992) |
| *Crepis vesicaria* | Asteraceae | Wild | Algeria | 1994 | 98.70 | Louadi and Doumandji (1998) |
| *Cucumis melo* | Cucurbitaceae | Cultivated | Italy | 1980 | 19.59 | Pinzauti (1981) |
| *Cucumis melo* | Cucurbitaceae | Cultivated | Spain | 2012 | 96.10 | Rodrigo Gómez et al. (2016) |
| *Cucumis sativus* | Cucurbitaceae | Cultivated | Algeria | 2001 | 66.20 | Benachour and Louadi (2011) |
| *Cucumis sativus* | Cucurbitaceae | Cultivated | Algeria | 2002 | 75.90 | Benachour and Louadi (2011) |
| *Cucurbita pepo* | Cucurbitaceae | Cultivated | Algeria | 2001 | 15.00 | Benachour (2008) |
| *Cucurbita pepo* | Cucurbitaceae | Cultivated | Algeria | 2003 | 0.00 | Benachour (2008) |
| *Cucurbita pepo* | Cucurbitaceae | Cultivated | Jordan | 2007 | 0.00 | Abu-Hammour and Wittmann ( 2010) |
| *Cytisus multiflorus* | Fabaceae | Wild | Spain | 1998 | 0.00 | Rodríguez-Riaño et al. (2004) |
| *Cytisus scoparius* | Fabaceae | Wild | Spain | 1996 | 0.00 | Malo and Baonza (2002) |
| *Cytisus scoparius* | Fabaceae | Wild | Spain | 2012 | 10.53 | Magrach et al. (2017) |
| *Cytisus striatus* | Fabaceae | Wild | Spain | 1995.5 | 2.00 | Rodríguez-Riaño et al. (1999) |
| *Daphne gnidium* | Thymelaeaceae | Wild | Spain | 1982.5 | 100.00 | J. Herrera (1985, 1988) |
| *Daucus carota* | Apiaceae | Cultivated | Egypt | 1981 | 51.10 | Hussein and Abdel-Aal (1982) |
| *Daucus carota* | Apiaceae | Cultivated | Italy | 1984 | 99.77 | Ricciardelli d'Albore (1986) |
| *Delphinium bolosii* | Ranunculaceae | Wild | Spain | 2002 | 94.19 | Orellana et al. (2008) |
| *Delphinium bolosii* | Ranunculaceae | Wild | Spain | 2002 | 97.98 | Orellana et al. (2008) |
| *Dictamnus albus* | Rutaceae | Wild | Italy | 2012.5 | 91.10 | Sgolastra et al. (2016) |
| *Digitalis lanata* | Plantaginaceae | Wild | Italy | 1983 | 93.00 | Ricciardelli d'Albore (1984) |
| *Digitalis purpurea* | Plantaginaceae | Wild | Italy | 1983 | 46.32 | Ricciardelli d'Albore (1984) |
| *Diplotaxis sp.* | Brassicaceae | Wild | Spain | 2014 | 42.51 | Pérez-Marcos et al. (2017) |
| *Drosophyllum lusitanicum* | Drosophyllaceae | Wild | Spain | 1994 | 100.00 | Ortega Olivencia et al. (1995) |
| *Ecballium elaterium* | Cucurbitaceae | Wild | France | 1996.5 | 84.00 | Rust et al. (2003) |
| *Echium plantagineum* | Boraginaceae | Wild | Spain | 1992 | 97.65 | Guitián et al. (1993) |
| *Echium plantagineum* | Boraginaceae | Wild | Spain | 2011 | 57.69 | Magrach et al. (2017) |
| *Echium vulgare* | Boraginaceae | Wild | Spain | 2014 | 92.93 | Pérez-Marcos et al. (2017) |
| *Elaeoselinum foetidum* | Apiaceae | Wild | Spain | 2012 | 61.29 | Magrach et al. (2017) |
| *Erica ciliaris* | Ericaceae | Wild | Spain | 1982.5 | 100.00 | J. Herrera (1985, 1988) |
| *Eriobotrya japonica* | Rosaceae | Cultivated | Jordan | 2005 | 15.00 | Freihat et al. (2008) |
| *Foeniculum vulgare* | Apiaceae | Cultivated | Italy | 1984 | 96.84 | Ricciardelli d'Albore (1986) |
| *Fragaria x ananassa* | Rosaceae | Cultivated | Portugal | 2003 | 50.57 | Albano et al. (2009) |
| *Fumana hispidula* | Cistaceae | Wild | Spain | 2009.5 | 95.60 | Carrió and Güemes (2013) |
| *Galactites tomentosa* | Asteraceae | Wild | Spain | 1986 | 33.01 | Bosch et al. (1997) |
| *Galactites tomentosa* | Asteraceae | Wild | Spain | 2011 | 17.07 | Magrach et al. (2017) |
| *Genista hirsuta* | Fabaceae | Wild | Spain | 2011 | 20.55 | Magrach et al. (2017) |
| *Genista hirsuta* | Fabaceae | Wild | Spain | 2012 | 40.86 | Magrach et al. (2017) |
| *Halimium commutatum* | Cistaceae | Wild | Spain | 1982.5 | 57.47 | J. Herrera (1985, 1988) |
| *Halimium halimifolium* | Cistaceae | Wild | Spain | 1982.5 | 71.43 | J. Herrera (1985, 1988) |
| *Halimium halimifolium* | Cistaceae | Wild | Spain | 2012 | 40.43 | Magrach et al. (2017) |
| *Hedera helix* | Araliaceae | Wild | Italy | 2003 | 0.33 | Vezza et al. (2006) |
| *Hedysarum coronarium* | Fabaceae | Cultivated | Italy | 1979 | 9.28 | Pinzauti and Magnani (1981) |
| *Hedysarum coronarium* | Fabaceae | Cultivated | Italy | 1983 | 5.17 | Ricciardelli d'Albore (1983) |
| *Hedysarum coronarium* | Fabaceae | Cultivated | Tunisia | 1983 | 57.38 | Sonet and Jacob-Remacle (1987) |
| *Hedysarum coronarium* | Fabaceae | Cultivated | Italy | 1998 | 27.32 | Acciaro et al. (2000) |
| *Hedysarum coronarium* | Fabaceae | Wild | Spain | 2009 | 23.50 | Montero-Castaño et al. (2014) |
| *Hedysarum coronarium* | Fabaceae | Wild | Spain | 2009 | 22.70 | Montero-Castaño et al. (2014) |
| *Hedysarum coronarium* | Fabaceae | Wild | Spain | 2010 | 12.50 | Montero-Castaño et al. (2014) |
| *Hedysarum coronarium* | Fabaceae | Wild | Spain | 2010 | 30.20 | Montero-Castaño et al. (2014) |
| *Helianthemum caput-felis* | Cistaceae | Wild | Spain | 2001 | 5.30 | Rodríguez-Pérez (2005) |
| *Helianthemum caput-felis* | Cistaceae | Wild | Spain | 2012 | 39.40 | Agulló et al. (2015) |
| *Helianthemum caput-felis* | Cistaceae | Wild | Spain | 2014 | 31.50 | Agulló et al. (2015) |
| *Helianthemum marifolium* | Cistaceae | Wild | Spain | 2001 | 4.40 | Rodríguez-Pérez (2005) |
| *Helianthus annuus* | Asteraceae | Cultivated | Egypt | 1981 | 3.80 | Hussein and Abdel-Aal (1982) |
| *Helianthus annuus* | Asteraceae | Cultivated | Spain | 1987 | 35.09 | Ortiz-Sánchez and Tinaut (1994) |
| *Helianthus annuus* | Asteraceae | Cultivated | Turkey | 1995.5 | 16.00 | Calmazur and Ozbek (1999) |
| *Helianthus annuus* | Asteraceae | Cultivated | Turkey | 2006 | 6.89 | Aytekin and Cagatay (2008) |
| *Helianthus annuus* | Asteraceae | Cultivated | Israel | 2010.5 | 6.60 | Pisanty et al. (2014) |
| *Helianthus annuus* | Asteraceae | Cultivated | Spain | 2013 | 4.37 | Hevia et al. (2016) |
| *Helianthus annuus* | Asteraceae | Cultivated | Italy | 2014 | 2.21 | Bartual et al. (2018) |
| *Helianthus annuus* | Asteraceae | Cultivated | Italy | 2015 | 6.11 | Bartual et al. (2018) |
| *Helichrysum picardii* | Asteraceae | Wild | Spain | 1982.5 | 100.00 | J. Herrera (1985, 1988) |
| *Helleborus foetidus* | Ranunculaceae | Wild | Spain | 1998 | 100.00 | C. M. Herrera et al. (2001) |
| *Helleborus foetidus* | Ranunculaceae | Wild | Spain | 1998 | 100.00 | C. M. Herrera et al. (2001) |
| *Helleborus foetidus* | Ranunculaceae | Wild | Spain | 1998 | 88.06 | C. M. Herrera et al. (2001) |
| *Helleborus foetidus* | Ranunculaceae | Wild | Spain | 1999 | 100.00 | C. M. Herrera et al. (2001) |
| *Helleborus foetidus* | Ranunculaceae | Wild | Spain | 1999 | 100.00 | C. M. Herrera et al. (2001) |
| *Helleborus foetidus* | Ranunculaceae | Wild | Spain | 1999 | 78.58 | C. M. Herrera et al. (2001) |
| *Hibiscus sabdariffa* | Malvaceae | Cultivated | Egypt | 2006 | 0.00 | Abdel-Moniem et al. (2011) |
| *Hibiscus sabdariffa* | Malvaceae | Cultivated | Egypt | 2007 | 0.00 | Abdel-Moniem et al. (2011) |
| *Iris cedretii* | Iridaceae | Wild | Lebanon | 2004 | 96.00 | Monty (2004) |
| *Iris sofarana* | Iridaceae | Wild | Lebanon | 2004 | 100.00 | Monty (2004) |
| *Jankaea heldreichii* | Gesneriaceae | Wild | Greece | 1987 | 100.00 | Vokou et. Al. (1990) |
| *Jasminum fruticans* | Oleaceae | Wild | Spain | 1994.5 | 99.30 | Guitián et al. (1998) |
| *Jatropha curcas* | Euphorbiaceae | Wild | Israel | 2009 | 12.98 | Samra et al. (2014) |
| *Lathyrus clymenum* | Fabaceae | Wild | Spain | 1986 | 9.10 | Bosch et al. (1997) |
| *Lathyrus latifolius* | Fabaceae | Wild | Spain | 1986 | 7.53 | Bosch et al. (1997) |
| *Lavandula latifolia* | Lamiaceae | Wild | Spain | 1982 | 55.40 | C. M. Herrera (1988) |
| *Lavandula latifolia* | Lamiaceae | Wild | Spain | 1983 | 40.23 | C. M. Herrera (1988) |
| *Lavandula latifolia* | Lamiaceae | Wild | Spain | 1984 | 58.55 | C. M. Herrera (1988) |
| *Lavandula latifolia* | Lamiaceae | Wild | Spain | 1985 | 66.22 | C. M. Herrera (1988) |
| *Lavandula latifolia* | Lamiaceae | Wild | Spain | 1986 | 48.04 | C. M. Herrera (1988) |
| *Lavandula latifolia* | Lamiaceae | Wild | Spain | 1987 | 33.29 | C. M. Herrera (1988) |
| *Lavandula latifolia* | Lamiaceae | Wild | Spain | 1991 | 42.55 | C. M. Herrera (2000) |
| *Lavandula officinalis* | Lamiaceae | Cultivated | Algeria | 2009 | 40.10 | Benachour (2017) |
| *Lavandula officinalis* | Lamiaceae | Cultivated | Algeria | 2010 | 28.40 | Benachour (2017) |
| *Lavandula officinalis* | Lamiaceae | Cultivated | Algeria | 2013 | 33.60 | Benachour (2017) |
| *Lavandula stoechas* | Lamiaceae | Wild | Spain | 1982.5 | 3.90 | J. Herrera (1985, 1988) |
| *Lavandula stoechas* | Lamiaceae | Wild | Spain | 2011 | 13.30 | Magrach et al. (2017) |
| *Lavandula stoechas* | Lamiaceae | Wild | Spain | 2012 | 27.28 | Magrach et al. (2017) |
| *Leontodon longirostris* | Asteraceae | Wild | Spain | 2011 | 99.28 | Magrach et al. (2017) |
| *Linaria lilacina* | Plantaginaceae | Wild | Spain | 2005 | 38.64 | Sánchez-Lafuente (2007) |
| *Lobularia maritima* | Brassicaceae | Wild | Spain | 1986 | 90.54 | Bosch et al. (1997) |
| *Lonicera arborea* | Caprifoliaceae | Wild | Spain | 1989 | 71.52 | Jordano (1990) |
| *Lonicera periclymenum* | Caprifoliaceae | Wild | Spain | 1982.5 | 100.00 | J. Herrera (1985, 1988) |
| *Lotus corniculatus* | Fabaceae | Cultivated | Italy | 1983 | 83.84 | Ricciardelli d'Albore (1984) |
| *Lupinus albus* | Fabaceae | Cultivated | Italy | 1983 | 88.38 | Ricciardelli d'Albore (1983) |
| *Lythrum salicaria* | Lythraceae | Wild | Italy | 2014 | 49.20 | Benvenuti et al. (2016) |
| *Malus domestica* | Rosaceae | Cultivated | Italy | 1963 | 27.66 | Solinas and Bin (1964) |
| *Malus domestica* | Rosaceae | Cultivated | Italy | 1964 | 21.62 | Solinas and Bin (1964) |
| *Malus domestica* | Rosaceae | Cultivated | Italy | 1964 | 94.20 | Solinas and Bin (1964) |
| *Malus domestica* | Rosaceae | Cultivated | Spain | 1993 | 40.20 | Vicens and Bosch (2000) |
| *Malus domestica* | Rosaceae | Cultivated | Spain | 1994 | 11.90 | Vicens and Bosch (2000) |
| *Malus domestica* | Rosaceae | Cultivated | Spain | 1995 | 35.80 | Vicens and Bosch (2000) |
| *Malus domestica* | Rosaceae | Cultivated | Italy | 2009 | 60.58 | Marini et al. (2012) |
| *Malva sylvestris* | Malvaceae | Wild | Algeria | 1994 | 97.44 | Louadi and Doumandji (1998) |
| *Mangifera indica* | Anacardiaceae | Cultivated | Israel | 1995 | 29.50 | Dag and Gazit (2000) |
| *Matricaria chamomilla* | Asteraceae | Cultivated | Egypt | 1981 | 76.90 | Hussein and Abdel-Aal (1982) |
| *Medicago arborea* | Fabaceae | Cultivated | Italy | 1983 | 19.39 | Ricciardelli d'Albore (1984) |
| *Medicago sativa* | Fabaceae | Cultivated | Turkey | 1973 | 99.46 | Özbek (1976) |
| *Medicago sativa* | Fabaceae | Cultivated | Italy | 1983 | 63.90 | Ricciardelli d'Albore (1984) |
| *Moricandia nitens* | Brassicaceae | Wild | Israel | 1989 | 100.00 | Küchmeister et al. (1995) |
| *Muscari comosum* | Hyacinthaceae | Wild | Italy | 2010 | 90.50 | Canale et al. (2014) |
| *Myrtus communis* | Myrtaceae | Wild | Spain | 1982.5 | 100.00 | J. Herrera (1985, 1988) |
| *Myrtus communis* | Myrtaceae | Wild | Spain | 2007 | 53.40 | González-Varo et al. (2009) |
| *Myrtus communis* | Myrtaceae | Wild | Spain | 2007 | 78.20 | González-Varo et al. (2009) |
| *Myrtus communis* | Myrtaceae | Wild | Spain | 2007 | 79.50 | González-Varo et al. (2009) |
| *Myrtus communis* | Myrtaceae | Wild | Spain | 2007 | 3.90 | González-Varo et al. (2009) |
| *Myrtus communis* | Myrtaceae | Wild | Spain | 2007 | 1.20 | González-Varo et al. (2009) |
| *Myrtus communis* | Myrtaceae | Wild | Spain | 2007 | 49.40 | González-Varo et al. (2009) |
| *Narcissus albimarginatus* | Amaryllidaceae | Wild | Morocco | 2004 | 79.25 | Pérez-Barrales et al. (2006) |
| *Narcissus broussonetii* | Amaryllidaceae | Wild | Morocco | 2009 | 70.47 | Santos-Gally et al. (2015) |
| *Narcissus cavanillesii* | Amaryllidaceae | Wild | Portugal | 2002 | 100.00 | Marques et al. (2007) |
| *Narcissus serotinus* | Amaryllidaceae | Wild | Spain | 1996.5 | 81.05 | Pérez Chiscano (2016) |
| *Narcissus serotinus* | Amaryllidaceae | Wild | Portugal | 2002 | 100.00 | Marques et al. (2007) |
| *Narcissus tazetta* | Amaryllidaceae | Wild | Israel | 1987 | 100.00 | Arroyo and Dafni (1995) |
| *Narcissus tazetta* | Amaryllidaceae | Wild | Israel | 1988 | 0.00 | Arroyo and Dafni (1995) |
| *Onobrychis viciifolia* | Fabaceae | Cultivated | Italy | 1983 | 4.04 | Ricciardelli d'Albore (1984) |
| *Ononis natrix* | Fabaceae | Wild | Spain | 1986 | 100.00 | Bosch et al. (1997) |
| *Orchis coriophora* | Orchidaceae | Wild | Israel | 1973 | 50.00 | Dafni and Ivri (1979) |
| *Origanum majorana* | Lamiaceae | Wild | Italy | 1982 | 74.01 | Ricciardelli d'Albore (1983) |
| *Origanum syriacum* | Lamiaceae | Cultivated | Jordan | 2006.5 | 64.20 | Al-Ghzawi et al. (2009) |
| *Origanum vulgare* | Lamiaceae | Wild | Italy | 1982 | 23.80 | Ricciardelli d'Albore (1983) |
| *Osyris lanceolata* | Santalaceae | Wild | Spain | 1982.5 | 53.85 | J. Herrera (1985, 1988) |
| *Oxalis pes-caprae* | Oxalidaceae | Wild | Algeria | 1994 | 83.33 | Louadi and Doumandji (1998) |
| *Paeonia broteroi* | Paeoniaceae | Wild | Spain | 1997 | 63.90 | Sánchez-Lafuente (2002) |
| *Paeonia broteroi* | Paeoniaceae | Wild | Spain | 1997 | 71.40 | Sánchez-Lafuente (2002) |
| *Paeonia broteroi* | Paeoniaceae | Wild | Spain | 1998 | 59.20 | Sánchez-Lafuente (2002) |
| *Paeonia broteroi* | Paeoniaceae | Wild | Spain | 1998 | 68.00 | Sánchez-Lafuente (2002) |
| *Pancratium parviflorum* | Amaryllidaceae | Wild | Israel | 1988 | 100.00 | Dafni (1996) |
| *Papaver rhoeas* | Papaveraceae | Wild | Spain | 1986 | 66.67 | Bosch et al. (1997) |
| *Papaver rhoeas* | Papaveraceae | Wild | Algeria | 1994 | 90.53 | Louadi and Doumandji (1998) |
| *Petrocoptis grandiflora* | Caryophyllaceae | Wild | Spain | 1992 | 100.00 | Navarro et al. (1993) |
| *Petrocoptis grandiflora* | Caryophyllaceae | Wild | Spain | 1993 | 100.00 | Guitián et al. (1994) |
| *Petrocoptis montsicciana* | Caryophyllaceae | Wild | Spain | 1999 | 98.80 | Bosch et al. (2002) |
| *Petroselinum crispum* | Apiaceae | Cultivated | Italy | 1984 | 99.73 | Ricciardelli d'Albore (1986) |
| *Pimpinella anisum* | Apiaceae | Cultivated | Italy | 1984 | 97.54 | Ricciardelli d'Albore (1986) |
| *Pimpinella anisum* | Apiaceae | Cultivated | Egypt | 2008 | 24.90 | Abd El-Wahab et al. (2012) |
| *Pimpinella anisum* | Apiaceae | Cultivated | Egypt | 2009 | 26.70 | Abd El-Wahab et al. (2012) |
| *Pisum sativum* | Fabaceae | Cultivated | Algeria | 2001 | 98.95 | Benachour (2008) |
| *Prunus dulcis* | Rosaceae | Cultivated | Italy | 1975 | 76.32 | Moleas (1978) |
| *Prunus dulcis* | Rosaceae | Cultivated | Italy | 1976 | 71.23 | Moleas (1978) |
| *Prunus dulcis* | Rosaceae | Cultivated | Italy | 1977 | 15.00 | Moleas (1978) |
| *Prunus dulcis* | Rosaceae | Cultivated | Italy | 1978 | 57.76 | Moleas (1978) |
| *Prunus dulcis* | Rosaceae | Cultivated | Spain | 1988 | 1.47 | Ortiz-Sánchez and Tinaut-Ranera (1995) |
| *Prunus dulcis* | Rosaceae | Cultivated | Israel | 2008 | 5.00 | Mandelik and Roll (2009) |
| *Prunus dulcis* | Rosaceae | Cultivated | Egypt | 2014 | 13.20 | Norfolk et al. (2016) |
| *Prunus dulcis* | Rosaceae | Cultivated | Spain | 2015.5 | 4.79 | Alomar et al. (2018) |
| *Prunus mahaleb* | Rosaceae | Wild | Spain | 1989 | 97.21 | Jordano (1993) |
| *Prunus mahaleb* | Rosaceae | Wild | Spain | 1990 | 16.30 | Guitián et al. (1993) |
| *Prunus salicina* | Rosaceae | Cultivated | Algeria | 2009 | 0.00 | Benachour and Louadi (2013) |
| *Prunus salicina* | Rosaceae | Cultivated | Algeria | 2010 | 0.80 | Benachour and Louadi (2013) |
| *Prunus spinosa* | Rosaceae | Wild | Spain | 1990 | 29.10 | Guitián et al. (1993) |
| *Psoralea bituminosa* | Fabaceae | Wild | Spain | 1986 | 73.33 | Bosch et al. (1997) |
| *Putoria calabrica* | Rubiaceae | Wild | Spain | 1996 | 71.43 | Ortiz et al. (2000) |
| *Pyrus communis* | Rosaceae | Cultivated | Italy | 1963 | 24.54 | Solinas and Bin (1964) |
| *Pyrus communis* | Rosaceae | Cultivated | Italy | 1964 | 13.21 | Solinas and Bin (1964) |
| *Pyrus communis* | Rosaceae | Cultivated | Italy | 1964 | 95.04 | Solinas and Bin (1964) |
| *Pyrus communis* | Rosaceae | Cultivated | Spain | 1996 | 15.70 | Monzón et al. (2004) |
| *Raphanus sativus* | Brassicaceae | Cultivated | Egypt | 1981 | 85.00 | Hussein and Abdel-Aal (1982) |
| *Raphanus sativus* | Brassicaceae | Cultivated | Algeria | 2004 | 96.88 | Benachour (2008) |
| *Reichardia picroides* | Asteraceae | Wild | Spain | 1986 | 100.00 | Bosch et al. (1997) |
| *Retama sphaerocarpa* | Fabaceae | Wild | Spain | 1995.5 | 0.00 | Rodríguez-Riaño et al. (1999) |
| *Rhamnus alaternus* | Rhamnaceae | Wild | Italy | 2014 | 12.20 | Canale et al. (2016) |
| *Rhododendron ponticum* | Ericaceae | Wild | Spain | 2002 | 100.00 | Stout et al. (2006) |
| *Rhus coriaria* | Anacardiaceae | Wild | Jordan | 2005 | 49.30 | Zaitoun et al. (2007) |
| *Rosmarinus officinalis* | Lamiaceae | Wild | Italy | 1982 | 7.50 | Ricciardelli d'Albore (1983) |
| *Rosmarinus officinalis* | Lamiaceae | Wild | Spain | 1982.5 | 23.88 | J. Herrera (1985, 1988) |
| *Rosmarinus officinalis* | Lamiaceae | Wild | Algeria | 1994 | 58.41 | Louadi and Doumandji (1998) |
| *Rosmarinus officinalis* | Lamiaceae | Wild | Israel | 2002 | 42.90 | Keasar et al. (2008) |
| *Rosmarinus officinalis* | Lamiaceae | Wild | Spain | 2011 | 18.37 | Magrach et al. (2017) |
| *Rosmarinus officinalis* | Lamiaceae | Wild | Spain | 2012 | 20.63 | Magrach et al. (2017) |
| *Rubus fruticosus* | Rosaceae | Wild | Italy | 1990 | 3.30 | Serini (1992) |
| *Rubus idaeus* | Rosaceae | Wild | Italy | 1990 | 14.14 | Serini (1992) |
| *Rubus ulmifolius* | Rosaceae | Wild | Spain | 1982.5 | 98.60 | J. Herrera (1985, 1988) |
| *Salvia fruticosa* | Lamiaceae | Wild | Israel | 1995 | 100.00 | Ne'eman and Dafni (1999) |
| *Salvia officinalis* | Lamiaceae | Wild | Italy | 1982 | 33.58 | Ricciardelli d'Albore (1983) |
| *Salvia sclarea* | Lamiaceae | Wild | Italy | 1982 | 99.00 | Ricciardelli d'Albore (1983) |
| *Salvia verbenaca* | Lamiaceae | Wild | Spain | 2014 | 100.00 | Pérez-Marcos et al. (2017) |
| *Satureja thymbra* | Lamiaceae | Wild | Israel | 1997 | 90.40 | Potts et al. (2001) |
| *Satureja thymbra* | Lamiaceae | Wild | Israel | 1997 | 82.10 | Potts et al. (2001) |
| *Scabiosa atropurpurea* | Dipsacaceae | Wild | Spain | 1986 | 30.13 | Bosch et al. (1997) |
| *Scolymus hispanicus* | Asteraceae | Wild | Algeria | 1994 | 100.00 | Louadi and Doumandji (1998) |
| *Scrophularia grandiflora* | Scrophulariaceae | Wild | Portugal | 2009 | 80.10 | Ortega-Olivencia et al. (2012) |
| *Scrophularia grandiflora* | Scrophulariaceae | Wild | Portugal | 2009 | 99.20 | Ortega-Olivencia et al. (2012) |
| *Scrophularia grandiflora* | Scrophulariaceae | Wild | Portugal | 2010 | 87.20 | Ortega-Olivencia et al. (2012) |
| *Scrophularia grandiflora* | Scrophulariaceae | Wild | Portugal | 2010 | 96.40 | Ortega-Olivencia et al. (2012) |
| *Scrophularia sambucifolia* | Scrophulariaceae | Wild | Spain | 2009 | 39.90 | Ortega-Olivencia et al. (2012) |
| *Scrophularia sambucifolia* | Scrophulariaceae | Wild | Spain | 2009 | 4.70 | Ortega-Olivencia et al. (2012) |
| *Scrophularia sambucifolia* | Scrophulariaceae | Wild | Spain | 2010 | 30.30 | Ortega-Olivencia et al. (2012) |
| *Scrophularia sambucifolia* | Scrophulariaceae | Wild | Spain | 2010 | 1.60 | Ortega-Olivencia et al. (2012) |
| *Sedum sediforme* | Crassulaceae | Wild | Spain | 1986 | 72.38 | Bosch et al. (1997) |
| *Senecio nebrodensis* | Asteraceae | Wild | Algeria | 1994 | 50.00 | Louadi and Doumandji (1998) |
| *Sesamum indicum* | Pedaliaceae | Cultivated | Egypt | 2010 | 44.80 | Mahmoud (2012) |
| *Sesamum indicum* | Pedaliaceae | Cultivated | Egypt | 2011 | 40.00 | Mahmoud (2012) |
| *Sesamum indicum* | Pedaliaceae | Cultivated | Egypt | 2011 | 77.30 | Kamel et al. (2013) |
| *Sesamum indicum* | Pedaliaceae | Cultivated | Egypt | 2012 | 81.90 | Kamel et al. (2013) |
| *Seseli farrenyi* | Apiaceae | Wild | Spain | 1999 | 95.40 | Rovira et al. (2002) |
| *Silene acutifolia* | Caryophyllaceae | Wild | Spain | 1997 | 99.73 | Buide (2006) |
| *Silene acutifolia* | Caryophyllaceae | Wild | Spain | 1998 | 99.89 | Buide (2006) |
| *Silene vulgaris* | Caryophyllaceae | Wild | Spain | 2014 | 97.22 | Pérez-Marcos et al. (2017) |
| *Sinapis arvensis* | Brassicaceae | Wild | Algeria | 1994 | 37.50 | Louadi and Doumandji (1998) |
| *Sonchus tenerrimus* | Asteraceae | Wild | Spain | 1986 | 80.86 | Bosch et al. (1997) |
| *Stauracanthus genistoides* | Fabaceae | Wild | Spain | 1982.5 | 43.75 | J. Herrera (1985, 1988) |
| *Sternbergia clusiana* | Amaryllidaceae | Wild | Israel | 1977.5 | 23.86 | Dafni and Werker (1982) |
| *Teucrium fruticans* | Lamiaceae | Wild | Spain | 2012 | 58.00 | Magrach et al. (2017) |
| *Thymus capitatus* | Lamiaceae | Wild | Greece | 2006 | 6.10 | Tscheulin and Petanidou (2011) |
| *Thymus longicaulis* | Lamiaceae | Wild | Italy | 2013 | 78.80 | Campolo et al. (2016) |
| *Thymus loscosii* | Lamiaceae | Wild | Spain | 2002 | 19.50 | Orellana et al. (2005) |
| *Thymus tomentosus* | Lamiaceae | Wild | Spain | 1982.5 | 100.00 | J. Herrera (1985, 1988) |
| *Thymus vulgaris* | Lamiaceae | Wild | Spain | 2008 | 74.75 | Arnan et al. (2014) |
| *Tolpis barbata* | Asteraceae | Wild | Spain | 2011 | 100.00 | Magrach et al. (2017) |
| *Tolpis barbata* | Asteraceae | Wild | Spain | 2012 | 100.00 | Magrach et al. (2017) |
| *Trifolium alexandrinum* | Fabaceae | Cultivated | Egypt | 1981 | 32.00 | Hussein and Abdel-Aal (1982) |
| *Trifolium pratense* | Fabaceae | Cultivated | Italy | 1983 | 89.95 | Ricciardelli d'Albore (1983) |
| *Trifolium repens* | Fabaceae | Cultivated | Italy | 1987 | 80.27 | Ortu et al. (1991) |
| *Ulex minor* | Fabaceae | Wild | Spain | 1982.5 | 4.44 | J. Herrera (1985, 1988) |
| *Ulex parviflorus* | Fabaceae | Wild | Spain | 1982.5 | 0.00 | J. Herrera (1985, 1988) |
| *Ulex parviflorus* | Fabaceae | Wild | Spain | 2014 | 7.02 | Castellanos et al. (2019) |
| *Urginea maritima* | Asparagaceae | Wild | Israel | 1983.5 | 85.29 | Dafni and Dukas (1986) |
| *Vaccinium corymbosum* | Ericaceae | Cultivated | Italy | 1990 | 69.32 | Serini (1992) |
| *Vaccinium corymbosum* | Ericaceae | Cultivated | Italy | 1998 | 84.32 | Prodorutti et al. (2005) |
| *Vaccinium corymbosum* | Ericaceae | Cultivated | Italy | 1999 | 93.22 | Prodorutti et al. (2005) |
| *Vaccinium corymbosum* | Ericaceae | Cultivated | Italy | 2000 | 94.99 | Prodorutti et al. (2005) |
| *Vicia cracca* | Fabaceae | Cultivated | Italy | 1983 | 59.99 | Ricciardelli d'Albore (1983) |
| *Vicia faba* | Fabaceae | Cultivated | France | 1973 | 20.90 | Tasei (1976) |
| *Vicia faba* | Fabaceae | Cultivated | Egypt | 1981 | 12.50 | Hussein and Abdel-Aal (1982) |
| *Vicia faba* | Fabaceae | Cultivated | France | 1994 | 30.80 | Pierre et al. (1996) |
| *Vicia faba* | Fabaceae | Cultivated | France | 1998 | 51.50 | Pierre et al. (1999) |
| *Vicia faba* | Fabaceae | Cultivated | Spain | 1998 | 96.10 | Pierre et al. (1999) |
| *Vicia faba* | Fabaceae | Cultivated | Algeria | 2000 | 75.00 | Benachour et al. (2007) |
| *Vicia faba* | Fabaceae | Cultivated | Algeria | 2001 | 60.00 | Benachour et al. (2007) |
| *Vicia faba* | Fabaceae | Cultivated | Algeria | 2002 | 94.12 | Benachour et al. (2007) |
| *Vicia faba* | Fabaceae | Cultivated | Algeria | 2003 | 57.78 | Aouar-Sadli et al. (2008) |
| *Vicia sativa* | Fabaceae | Wild | Spain | 2014 | 37.50 | Pérez-Marcos et al. (2017) |
| *Vicia villosa* | Fabaceae | Cultivated | Jordan | 2005 | 44.50 | Al-Ghzawi et al. (2009) |
| *Vitex agnus-castus* | Verbenaceae | Wild | Greece | 2008 | 58.65 | Barthell et al. (2012) |

*** References**

Abd El-Wahab, T. E., I. M. A. Ebadah, and Y. A. Mahmoud. 2012. Insect pollinators of anise plants (*Pimpinella anisum* L.) and the important role of honey bees (*Apis mellifera* L.) on their yield productivity. Archives of Phytopathology and Plant Protection 45:677-685.

Abdel-Moniem, A. S. H., T. E. Abd El-Wahab, and N. A. Farag. 2011. Prevailing insects in Roselle plants, *Hibiscus sabdariffa* L., and their efficiency on pollination. Archives of Phytopathology and Plant Protection 44:242-252.

Abu-Hammour, K. 2008. Pollination of medicinal plants (*Nigella sativa* and *Coriandrum sativum*) and *Cucurbita pepo* in Jordan. Dissertation. Institut für Nutzpflanzenwissenschaften und Ressoucenschutz, University of Bonn.

Abu-Hammour, K., and D. Wittmann. 2010. Pollination and pollinators of *Cucurbita pepo* (Cucurbitaceae) in the Jordan Valley to improve seed set. Advances in Horticultural Science 24:249-256.

Acciaro, M., I. Floris, A. Lentini, A. Satta, and L. Sulas. 2000. Insect pollination of sulla (*Hedysarum coronarium* L.) and its effect on seed production in a Mediterranean environment. Cahiers Options Méditerranéennes 45:373-377.

Agulló, J. C., C. Pérez-Bañón, M. B. Crespo, and A. Juan. 2015. Puzzling out the reproductive biology of the endangered cat's head rockrose (*Helianthemum caput-felis*, Cistaceae). Flora 217:75-81.

Al-Ghzawi, A. A.-M., S. Zaitoun, N. Freihat, and A. Alqudah. 2009. Effect of pollination on seed set of *Origanum syriacum* under semiarid Mediterranean conditions. Acta Agriculturae Scandinavica, Section B — Soil & Plant Science 59:273-278.

Albano, S., E. Salvado, P. A. V. Borges, and A. Mexia. 2009. Floral visitors, their frequency, activity rate and Index of Visitation Rate in the strawberry fields of Ribatejo, Portugal: selection of potential pollinators. Part 1. Advances in Horticultural Science:238-245.

Alomar, D., M. A. González-Estévez, A. Traveset, and A. Lázaro. 2018. The intertwined effects of natural vegetation, local flower community, and pollinator diversity on the production of almond trees. Agriculture, Ecosystems & Environment 264:34-43.

Aouar-Sadli, M., K. Louadi, and S.-E. Doum. 2008. Pollination of the broad bean (*Vicia faba* L. var. *major*) (Fabaceae) by wild bees and honey bees (Hymenoptera: Apoidea) and its impact on the seed production in the Tizi-Ouzou area (Algeria). African Journal of Agricultural Research 3:266-272.

Arnan, X., A. Escolà, A. Rodrigo, and J. Bosch. 2014. Female reproductive success in gynodioecious *Thymus vulgaris*: pollen versus nutrient limitation and pollinator foraging behaviour. Botanical Journal of the Linnean Society 175:395-408.

Arroyo, J., and A. Dafni. 1995. Variations in habitat, season, flower traits and pollinators in dimorphic *Narcissus tazetta* L. (Amaryllidaceae) in Israel. New Phytologist 129:135-145.

Aytekin, A. M., and N. Cagatay. 2008. Observations on the pollination of sunflower (*Helianthus annuus* L.). Mellifera 8:2-7.

Barthell, J. F., J. M. Hranitz, J. R. Redd, M. L. Clement, K. C. Crocker, E. C. Becker, K. D. Leavitt, B. M. McCall, M. Mills-Novoa, and C. M. Walker. 2012. Observations on nectar availability and bee visitation at patches of yellow star-thistle and chasteberry on the northeast Aegean Island of Lesvos (Greece). Uludağ Bee Journal 12:55-61.

Bartual, A. M., G. Bocci, S. Marini, and A. C. Moonen. 2018. Local and landscape factors affect sunflower pollination in a Mediterranean agroecosystem. PLoS One 13:e0203990.

Benachour, K. 2008. Diversité et activité pollinisatrice des abeilles (Hymenoptera: Apoidea) sur les plantes cultivées. Ph. D. Thesis. Faculté des Sciences de la Nature et de la Vie, Université Mentouri de Constantine, Constantine, Algeria.

Benachour, K. 2017. Insect visitors of lavender (*Lavandula officinalis* L.): comparison of quantitative and qualitative interactions of the plant with its main pollinators. African Entomology 25:435-444.

Benachour, K., and K. Louadi. 2011. Comportement de butinage des abeilles (Hymenoptera: Apoidea) sur les fleurs mâles et femelles du concombre (*Cucumis sativus* L.)(Cucurbitaceae) en région de Constantine (Algérie). Annales de la Société Entomologique de France 47:63-70.

Benachour, K., and K. Louadi. 2013. Inventory of insect visitors, foraging behaviour and pollination efficiency of honeybees (*Apis mellifera* L.) (Hymenoptera: Apidae) on plum (*Prunus salicina* Lindl.)(Rosaceae) in the Constantine area, Algeria. African Entomology 21:354-361.

Benachour, K., K. Louadi, and M. Terzo. 2007. Rôle des abeilles sauvages et domestiques (Hymenoptera: Apoidea) dans la pollinisation de la fève (*Vicia faba* L. var. *major*)(Fabaceae) en région de Constantine (Algérie). Annales de la Société Entomologique de France 43:213-219.

Bendifallah, L., K. Louadi, and S. Doumandji. 2013. Bee fauna potential visitors of coriander flowers *Coriandrum sativum* L.(Apiaceae) in the Mitidja area (Algeria). Journal of Apicultural Science 57:59-70.

Benelli, G., S. Benvenuti, N. Desneux, and A. Canale. 2014. *Cephalaria transsylvanica*-based flower strips as potential food source for bees during dry periods in European Mediterranean basin countries. PLoS One 9:e93153.

Benelli, G., S. Benvenuti, P. L. Scaramozzino, and A. Canale. 2017. Food for honeybees? Pollinators and seed set of *Anthyllis barba-jovis* L. (Fabaceae) in arid coastal areas of the Mediterranean basin. Saudi Journal of Biological Sciences 24:1056-1060.

Benvenuti, S., G. Benelli, N. Desneux, and A. Canale. 2016. Long lasting summer flowerings of *Lythrum salicaria* as honeybee-friendly flower spots in Mediterranean basin agricultural wetlands. Aquatic Botany 131:1-6.

Blionis, G. J., and D. Vokou. 2001. Pollination ecology of *Campanula* species on Mt Olympos, Greece. Ecography 24:287-297.

Bosch, J. 1992. Floral biology and pollinators of three co-occurring *Cistus* species (Cistaceae). Botanical Journal of the Linnean Society 109:39-55.

Bosch, M., J. Simon, C. Blanché, and J. Molerò. 1997. Pollination ecology in the Tribe Delphineae (Ranunculaceae) in W mediterranean area: Floral visitors and pollinator behaviour. Lagascalia 19:545-562.

Bosch, M., J. Simon, A. M. Rovira, J. Molero, and C. Blanché. 2002. Pollination ecology of the pre-Pyrenean endemic *Petrocoptis montsicciana* (Caryophyllaceae): effects of population size. Biological Journal of the Linnean Society 76:79-90.

Buide, M. L. 2006. Pollination ecology of *Silene acutifolia* (Caryophyllaceae): floral traits variation and pollinator attraction. Annals of Botany 97:289-297.

Calmasur, O., and H. Ozbek. 1999. Pollinator bees (Hymenoptera, Apoidea) on sunflower (*Helianthus annuus* L.) and their effects on seed setting in Erzurum region. Turkish Journal of Biology 23:73-89.

Campolo, O., L. Zappalà, A. Malacrinò, F. Laudani, and V. Palmeri. 2016. Bees visiting flowers of *Thymus longicaulis* (Lamiaceae). Plant Biosystems 150:1182-1188.

Canale, A., G. Benelli, and S. Benvenuti. 2014. First record of insect pollinators visiting *Muscari comosum* (L.) Miller (Liliaceae-Hyacinthaceae), an ancient Mediterranean food plant. Plant Biosystems 148:889-894.

Canale, A., S. Benvenuti, A. Raspi, and G. Benelli. 2016. Insect pollinators of the late winter flowering *Rhamnus alaternus* L., a candidate for honeybee-friendly scrubland spots in intensively managed agricultural areas. Plant 150:611-615.

Carrió, E., and J. Güemes. 2013. The role of a mixed mating system in the reproduction of a Mediterranean subshrub (*Fumana hispidula*, Cistaceae). Journal of Plant Research 126:33-40.

Castellanos, M. C., J. Montero-Pau, P. Ziarsolo, J. M. Blanca, J. Cañizares, and J. G. Pausas. 2019. A stable pollination environment limits current but not potential evolution of floral traits. bioRxiv:581827. Doi: 10.1101/581827

Dafni, A. 1996. Autumnal and winter pollination adaptations under Mediterranean conditions. Bocconea 5:171-181.

Dafni, A., and R. Dukas. 1986. Insect and wind pollination in *Urginea maritima* (Liliaceae). Plant Systematics and Evolution 154:1-10.

Dafni, A., and Y. Ivri. 1979. Pollination ecology of, and hybridization between*, Orchis coriophora* L. and *O. collina* Sol. ex Russ. (Orchidaceae) in Israel. New Phytologist 83:181-187.

Dafni, A., and Y. Ivri. 1981. The flower biology of *Cephalanthera longifolia* (Orchidaceae) — pollen imitation and facultative floral mimicry. Plant Systematics and Evolution 137:229-240.

Dafni, A., and E. Werker. 1982. Pollination ecology of *Sternbergia clusiana* (Ker-Gawler) Spreng. (Amaryllidaceae). New Phytologist 91:571-577.

Dafni, A., D. Eisikowitch, and Y. Ivri. 1987. Nectar flow and pollinators' efficiency in two co-occurring species of *Capparis* (Capparaceae) in Israel. Plant Systematics and Evolution 157:181-186.

Dag, A., and S. Gazit. 2000. Mango pollinators in Israel. Journal of Applied Horticulture 2:39-43.

Descamps, C., L. Moquet, M. Migon, and A.-L. Jacquemart. 2015. Diversity of the insect visitors on *Calluna vulgaris* (Ericaceae) in southern France heathlands. Journal of Insect Science 15.

Fijen, T. P. M., and D. Kleijn. 2017. How to efficiently obtain accurate estimates of flower visitation rates by pollinators. Basic and Applied Ecology 19:11-18.

Frediani, D., and M. Pinzauti. 1985. Osservazioni sull'impollinazione entomofila del cartamo (*Carthamus tinctorius* L.). Sementi Elette 30:13-16.

Freihat, N. M., A. A.-M. Al-Ghzawi, S. Zaitoun, and A. Alqudah. 2008. Fruit set and quality of loquats (*Eriobotrya japonica*) as effected by pollinations under sub-humid Mediterranean. Scientia Horticulturae 117:58-62.

González-Varo, J. P., J. Arroyo, and A. Aparicio. 2009. Effects of fragmentation on pollinator assemblage, pollen limitation and seed production of Mediterranean myrtle (*Myrtus communis*). Biological Conservation 142:1058-1065.

González-Varo, J. P., F. J. Ortiz-Sánchez, and M. Vilà. 2016. Total bee dependence on one flower species despite available congeners of similar floral shape. PLoS One 11:e0163122.

Goras, G., C. Tananaki, M. Dimou, T. Tscheulin, T. Petanidou, and A. Thrasyvoulou. 2016. Impact of honeybee (*Apis mellifera* L.) density on wild bee foraging behaviour. Journal of Apicultural Science 60:49-62.

Guitián, J., and M. Fuentes. 1992. Reproductive biology of *Crataegus monogyna* in northwestern Spain. Acta Oecologica 13:3-11.

Guitián, J., P. Guitián, and L. Navarro. 1993. Tamaño del núcleo de población y polinización en *Echium plantagineum*. Anales del Jardín Botánico de Madrid 51:65-72.

Guitián, J., J. M. Sánchez, and P. Guitián. 1994. Pollination ecology of *Petrocoptis grandiflora* Rothm. (Caryophyllaceae); a species endemic to the north-west part of the Iberian Peninsula. Botanical Journal of the Linnean Society 115:19-27.

Guitián, J., P. Guitián, and L. Navarro. 1996. Fruit set, fruit reduction, and fruiting strategy in *Cornus sanguinea* (Cornaceae). American Journal of Botany 83:744-748.

Guitián, J., P. Guitián, and M. Medrano. 1998. Floral biology of the distylous Mediterranean shrub *Jasminum fruticans* (Oleaceae). Nordic Journal of Botany 18:195-201.

Hevia, V., J. Bosch, F. M. Azcárate, E. Fernández, A. Rodrigo, H. Barril-Graells, and J. A. González. 2016. Bee diversity and abundance in a livestock drove road and its impact on pollination and seed set in adjacent sunflower fields. Agriculture Ecosystems & Environment 232:336-344.

Hussein, M. H., and S. A. Abdel-Aal. 1982. Wild and honey bees as pollinators of 10 plant species in Assiut area, Egypt. Zeitschrift für Angewandte Entomologie 93:342-346.

Herrera, C. M. 1988. Variation in mutualisms: the spatio-temporal mosaic of a pollinator assemblage. Biological Journal of the Linnean Society 35:95-125.

Herrera, C. M. 2000. Flower-to-seedling consequences of different pollination regimes in an insect-pollinated shrub. Ecology 81:15-29.

Herrera, C. M., A. M. Sánchez-Lafuente, M. Medrano, J. Guitián, X. Cerdá, and P. Rey. 2001, Geographical variation in autonomous self-pollination levels unrelated to pollinator service in *Helleborus foetidus* (Ranunculaceae). American Journal of Botany 88, 1025-1032.

Herrera, J. 1985. Biología reproductiva del matorral de Doñana. Ph. D. Thesis. Faculty of Biology, University of Sevilla, Sevilla, Spain.

Herrera, J. 1988. Pollination relationships in southern Spanish Mediterranean shrublands. Journal of Ecology 76:274-287.

Jordano, P. 1990. Biología de la reproducción de tres especies del género *Lonicera* (Caprifoliaceae) en la Sierra de Cazorla. Anales del Jardín Botánico de Madrid 48:31-52.

Jordano, P. 1993. Pollination biology of *Prunus mahaleb* L.: deferred consequences of gender variation for fecundity and seed size. Biological Journal of the Linnean Society 50:65-84.

Kamel, S. M., A. E. H. Blal, H. M. Mahfouz, and M. Said. 2013. Pollinator fauna of sesame crop (*Sesamum indicum* L.) in Ismailia Governorate, Egypt. Cercetări Agronomice în Moldova 46:53-64.

Kamel, S. M., H. M. Mahfouz, E.-F. A. H. Blal, M. Said, and M. F. Mahmoud. 2015. Diversity of insect pollinators with reference to their impact on yield production of canola (*Brassica napus* L.) in Ismailia, Egypt. Pesticidi i Fitomedicina 30:161-168.

Keasar, T., A. Sadeh, and A. Shmida. 2008. Variability in nectar production and standing crop, and their relation to pollinator visits in a Mediterranean shrub. Arthropod-Plant Interactions 2:117-123.

Küchmeister, H., A. Shmida, and G. Gottsberger. 1995. Phenology and pollination ecology of the desert plant *Moricandia nitens* (Brassicaceae) in the Negev, Israel. Advances in Geoecology 28:157-171.

Lázaro, A., and A. Traveset. 2005. Spatio-temporal variation in the pollination mode of *Buxus balearica* (Buxaceae), an ambophilous and selfing species: mainland-island comparison. Ecography 28:640-652.

Louadi, K., and S. Doumandji. 1998. Diversité et activité de butinage des abeilles (Hymenoptera: Apoidea) dans une pelouse à thérophytes de Constantine (Algérie). Canadian Entomologist 130:691-702.

Magrach, A., J. P. González-Varo, M. Boiffier, M. Vilà, and I. Bartomeus. 2017. Honeybee spillover reshuffles pollinator diets and affects plant reproductive success. Nature Ecology & Evolution 1:1299-1307.

Mahmoud, F. M. 2012. Insects associated with sesame (*Sesamum indicum* L.) and the impact of insect pollinators on crop production. Pesticidi i Fitomedicina 27:117-129.

Malo, J. E., and J. Baonza. 2002. Are there predictable clines in plant-pollinator interactions along altitudinal gradients? The example of *Cytisus scoparius* (L.) Link in the Sierra de Guadarrama (Central Spain). Diversity and Distributions 8:365-371.

Mandelik, Y., and U. Roll. 2009. Diversity patterns of wild bees in almond orchards and their surrounding landscape. Israel Journal of Plant Sciences 57:185-191.

Manetas, Y., and Y. Petropoulou. 2000. Nectar amount, pollinator visit duration and pollination success in the Mediterranean shrub *Cistus creticus*. Annals of Botany 86:815-820.

Marini, L., M. Quaranta, P. Fontana, J. C. Biesmeijer, and R. Bommarco. 2012. Landscape context and elevation affect pollinator communities in intensive apple orchards. Basic and Applied Ecology 13:681-689.

Marques, I., A. Rosselló-Graells, D. Draper, and J. M. Iriondo. 2007. Pollination ecology and hybridization between *Narcissus cavanillesii* A. Barra & G. Lopéz and *N. serotinus* L. in Portugal. Bocconea 21:65-75.

Miñarro, M., and K. W. Twizell. 2015. Pollination services provided by wild insects to kiwifruit (*Actinidia deliciosa*). Apidologie 46:276-285.

Moleas, T. 1978. Osservazioni sugli insetti impollinatori del mandorlo in terra di Bari. Annali della Facolta di Agraria, Bari 30:609-627.

Montero-Castaño, A., M. Vilà, and F. J. Ortiz-Sánchez. 2014. Pollination ecology of a plant in its native and introduced areas. Acta Oecologica-International Journal of Ecology 56:1-9.

Monty, A. 2004. Caractérisation écologique de la pollinisation des Iris Oncocycles endémiques du Liban. Degree Thesis. Faculté Universitaire des Sciences Agronomiques de Gembloux, Université de Liège, Belgium.

Monzón, V. H., J. Bosch, and J. Retana. 2004. Foraging behavior and pollinating effectiveness of *Osmia cornuta* (Hymenoptera: Megachilidae) and *Apis mellifera* (Hymenoptera: Apidae) on “Comice” pear. Apidologie 35:575-585.

Navarro, L. 2000. Pollination ecology of *Anthyllis vulneraria* subsp. *vulgaris* (Fabaceae): nectar robbers as pollinators. American Journal of Botany 87:980-985.

Navarro, L., J. Guitián, and P. Guitián. 1993. Reproductive biology of *Petrocoptis grandiflora* Rothm. (Caryophyllaceae), a species endemic to northwest Iberian Peninsula. Flora 188:253-261.

Ne’eman, G., and A. Dafni. 1999. Fire, bees, and seed production in a Mediterranean key species *Salvia fruticosa* Miller (Lamiaceae). Israel Journal of Plant Sciences 47:157-163.

Norfolk, O., M. P. Eichhorn, and F. Gilbert. 2016. Flowering ground vegetation benefits wild pollinators and fruit set of almond within arid smallholder orchards. Insect Conservation and Diversity 9:236-243.

Obeso, J. R. 1992. Pollination ecology and seed set in *Asphodelus albus* (Liliaceae) in northern Spain. Flora 187:219-226.

Orellana, M. R., A. M. Rovira, C. Blanché, and M. Bosch. 2005. Pollination and reproductive success in the gynodioecious endemic *Thymus loscosii* (Lamiaceae). Canadian Journal of Botany 83:183-193.

Orellana, M. R., A. M. Rovira, C. Blanché i Vergés, and M. Bosch. 2008. Effects of local abundance on pollination and reproduction in the narrow endemic endangered species *Delphinium bolosii* (Ranunculaceae). Orsis 23:27-46.

Ortega-Olivencia, A., T. Rodríguez-Riaño, J. L. Pérez-Bote, J. López, C. Mayo, F. J. Valtueña, and M. Navarro-Pérez. 2012. Insects, birds and lizards as pollinators of the largest-flowered *Scrophularia* of Europe and Macaronesia. Annals of Botany 109:153-167.

Ortega-Olivencia, A., J. P. Carrasco, and J. A. Devesa. 1995. Floral and reproductive biology of *Drosophyllum lusitanicum* (L.) Link (Droseraceae). Botanical Journal of the Linnean Society 118:331-351.

Ortiz-Sánchez, F. J., and A. Tinaut-Ranera. 1995. Insect pollination of almond (*Prunus dulcis* (Mill.)) in southern Spain and its effect on production. Agronomia Lusitana 45:289-295.

Ortiz-Sánchez, F. J., and A. Tinaut. 1994. Composición y dinámica de la comunidad de polinizadores potenciales del girasol (*Helianthus annuus* L.) en Granada (España). Boletín de Sanidad Vegetal – Plagas 20:737-756.

Ortiz, P. L., M. Arista, and S. Talavera. 2000. Pollination and breeding system of *Putoria calabrica* (Rubiaceae), a Mediterranean dwarf shrub. Plant Biology 2:325-330.

Ortu, S., I. Floris, and S. Pampaloni. 1991. Osservazioni su insetti impollinatori di Trifoglio bianco (*Trifolium repens* L.). Apicoltore Moderno 82:103-111.

Özbek, H. 1976. Pollinator bees on alfalfa in the Erzurum region of Turkey. Journal of Apicultural Research 15:145-148.

Pérez-Barrales, R., P. Vargas, and J. Arroyo. 2006. New evidence for the Darwinian hypothesis of heterostyly: breeding systems and pollinators in *Narcissus* sect. Apodanthi. New Phytologist 171:553-567.

Pérez-Marcos, M., F. J. Ortiz-Sánchez, E. López-Gallego, M. J. Ramírez-Soria, and J. A. Sanchez. 2017. The importance of the qualitative composition of floral margins to the maintenance of rich communities of bees. IOBC-WPRS Bulletin 122:83-87.

Pérez Chiscano, J. L. 2016. Datos sobre la reproducción de *Narcissus serotinus* Loefl. ex L. (Amaryllidaceae), en la comarca de La Serena, Extremadura, España. Folia Botanica Extremadurensis 9:35-40.

Pierre, J., J. LeGuen, M. H. P. Delègue, J. Mesquida, R. Marilleau, and G. Morin. 1996. Comparative study of nectar secretion and attractivity to bees of two lines of spring-type faba bean (*Vicia faba* L var *equina* Steudel). Apidologie 27:65-75.

Pierre, J., M. J. Suso, M. T. Moreno, R. Esnault, and J. Le Guen. 1999. Diversity and efficiency of the pollinating entomofauna (Hymenoptera : Apidae) of Faba bean (*Vicia faba* L.) in two locations in France and Spain. Annales de la Société Entomologique de France 35:312-318.

Pinzauti, M. 1981. L'Ape nell'impollinazione del melone. L'Agricoltura Italiana 110:447-457.

Pinzauti, M., and G. Magnani. 1981. Ricerche comparative sull'impollinazione della sulla (*Hedysarum coronarium* L.). L'Agricoltura Italiana 110:36-47.

Pisanty, G., A. M. Klein, and Y. Mandelik. 2014. Do wild bees complement honeybee pollination of confection sunflowers in Israel? Apidologie 45:235-247.

Potts, S. G., A. Dafni, and G. Ne’eman. 2001. Pollination of a core flowering shrub species in Mediterranean phrygana: variation in pollinator diversity, abundance and effectiveness in response to fire. Oikos 92:71-80.

Prodorutti, D., F. Frilli, and P. A. Belletti. 2005. L’impollinazione del mirtillo gigante americano. Notiziario ERSA 2-3: 34-36.

Quaranta, M., and G. C. Ricciardelli d'Albore. 1996. Gli insetti impollinatori di *Agastache foeniculum* Kuntze (Labiatae) in Umbria. Annali della Facoltà di Agraria di Perugia 47:287-295.

Ricciardelli d'Albore, G. C. 1983a. Osservazioni sugli insetti impollinatori di alcune Labiate di interesse erboristico (*Origanum majorana* L., *Origanum vulgare* L., *Rosmarinus officinalis* L., *Salvia officinalis* L. e *Salvia sclarea* L.) in un areale specializzato. Redia 66:283-293.

Ricciardelli d'Albore, G. C. 1983b. Osservazioni sugli insetti impollinatori di alcune Leguminose (*Trifolium pratense* L., *Vicia cracca* L., *Hedysarum coronarium* L., *Astragalus glycyphyllos* L., *Lupinus albus* L.) in un areale specializzato. Annali della Facoltà di Agraria di Perugia 37:149-160.

Ricciardelli d'Albore, G. C. 1984a. Osservazioni sugli insetti impollinatori di *Atropa belladonna* L., *Digitalis purpurea* L., *D. lanata* Ehrh., *Valeriana officinalis* L. in un areale specializzato. L'Apicoltore Moderno 75:165-172.

Ricciardelli d'Albore, G. C. 1984b. Osservazioni sugli insetti pronubi di alcune Leguminose (*Onobrychis viciifolia* Scop., *Lotus corniculatus* L., *Medicago arborea* L., *Medicago sativa* L.) in un areale specializzato. Redia 67:145-155.

Ricciardelli d'Albore, G. C. 1986. Les insectes pollinisateurs de quelques ombellifères d'intérêt agricole et condimentaire (*Angelica archangelica* L., *Carum carvi* L., *Petroselinum crispum* Aw Hill., *Apium graveolens* L., *Pimpinella anisum* L., *Daucus carota* L., *Foeniculum vulgare* Miller v. *azoricum* Thell.). Apidologie 17:107-124.

Rodrigo Gómez, S., C. Ornosa, J. Selfa, M. Guara, and C. Polidori. 2016. Small sweat bees (Hymenoptera: Halictidae) as potential major pollinators of melon (*Cucumis melo*) in the Mediterranean. Entomological Science 19:55-66.

Rodríguez-Pérez, J. 2005. Breeding system, flower visitors and seedling survival of two endangered species of *Helianthemum* (Cistaceae). Annals of Botany 95:1229-1236.

Rodríguez-Riaño, T., A. Ortega-Olivencia, and J. A. Devesa. 1999. Reproductive biology in two Genisteae (Papilionoideae) endemic of the western Mediterranean region: *Cytisus striatus* and *Retama sphaerocarpa*. Canadian Journal of Botany 77:809-820.

Rodríguez-Riaño, T., A. Ortega-Olivencia, and J. A. Devesa. 2004. Reproductive biology in *Cytisus multiflorus* (Fabaceae). Annales Botanici Fennici:179-188.

Rovira, A., M. Bosch, J. Molero, and C. Blanché. 2002. Pollination ecology and breeding system of the very narrow coastal endemic *Seseli farrenyi* (Apiaceae). Effects of population fragmentation. Nordic Journal of Botany 22:727-740.

Rust, R. W., B. E. Vaissière, and P. Westrich. 2003. Pollinator biodiversity and floral resource use in *Ecballium elaterium* (Cucurbitaceae), a Mediterranean endemic. Apidologie 34:29-42.

Samra, S., Y. Samocha, D. Eisikowitch, and Y. Vaknin. 2014. Can ants equal honeybees as effective pollinators of the energy crop *Jatropha curcas* L. under Mediterranean conditions? Global Change Biology – Bioenergy 6:756-767.

Sánchez-Lafuente, A. 2002. Floral variation in the generalist perennial herb *Paeonia broteroi* (Paeoniaceae): differences between regions with different pollinators and herbivores. American Journal of Botany 89:1260-1269.

Sánchez-Lafuente, A. M. 2007. Corolla herbivory, pollination success and fruit predation in complex flowers: an experimental study with *Linaria lilacina* (Scrophulariaceae). Annals of Botany 99:355-364.

Santos-Gally, R., A. de Castro, R. Pérez-Barrales, and J. Arroyo. 2015. Stylar polymorphism on the edge: unusual flower traits in Moroccan *Narcissus broussonetii* (Amaryllidaceae). Botanical Journal of the Linnean Society 177:644-656.

Schurr, L., L. Affre, F. Flacher, T. Tatoni, L. Le Mire Pecheux, and B. Geslin. 2019. Pollination insights for the conservation of a rare threatened plant species, *Astragalus tragacantha* (Fabaceae). Biodiversity and Conservation 28:1389-1409.

Serini, G. B. 1992. Seasonal variations in the pollinators *Apis mellifera* L. and *Bombus* spp. in the “Massiccio del Campo dei Fiori” area (Varese, Northern Italy). Ethology Ecology & Evolution 4:37-42.

Sgolastra, F., A. Fisogni, M. Quaranta, G. Bogo, L. Bortolotti, and M. Galloni. 2016. Temporal activity patterns in a flower visitor community of *Dictamnus albus* in relation to some biotic and abiotic factors. Bulletin of Insectology 69:291-300.

Solinas, M., and F. Bin. 1964. Osservazioni sull'attività degli insetti impollinatori del pero e del melo. Rivista della Ortoflorofrutticoltura Italiana 48:479-496.

Sonet, M., and A. Jacob-Remacle. 1987. Pollinisation de la légumineuse fourragère *Hedysarum coronarium* L. en Tunisie. Bulletin des Recherches Agronomiques de Gembloux 22:19-32.

Stout, J. C., J. A. N. Parnell, J. Arroyo, and T. P. Crowe. 2006. Pollination ecology and seed production of *Rhododendron ponticum* in native and exotic habitats. Biodiversity & Conservation 15:755-777.

Taha, E. A., and Y. A. Bayoumi. 2009. The value of honey bees (*Apis mellifera*, L.) as pollinators of summer seed watermelon (*Citrullus lanatus colothynthoides* L.) in Egypt. Acta Biologica Szegediensis 53:33-37.

Talavera, S., F. Bastida, P. L. Ortiz, and M. Arista. 2001. Pollinator attendance and reproductive success in *Cistus libanotis* L.(Cistaceae). International Journal of Plant Sciences 162:343-352.

Tasei, J. N. 1976. Les insectes pollinisateurs de la féverole d'hiver (*Vicia faba equina* L.) et la pollinisation des plantes mâle-stérile en production de semence hybride. Apidologie 7:1-28.

Torres, M. E., C. Ruiz, J. M. Iriondo, and C. Pérez. 2001. Pollination ecology of *Antirrhinum microphyllum* Rothm. (Scrophulariaceae). Bocconea 13:543-547.

Tscheulin, T., and T. Petanidou. 2011. Does spatial population structure affect seed set in pollen-limited *Thymus capitatus*? Apidologie 42:67-77.

Vanderbergh, M. 2002. Faune pollinisatrice des prés de fauche à centaurées (Asteraceae) de la Cerdagne (Pyrénées-Orientales, France). Rapport de recherche, Service de Zoologie, Université de Mons-Hainaut, Mons, Belgium.

Vezza, M., M. Nepi, M. Guarnieri, D. Artese, N. Rascio, and E. Pacini. 2006. Ivy (*Hedera helix* L.) flower nectar and nectary ecophysiology. International Journal of Plant Sciences 167:519-527.

Vicens, N., and J. Bosch. 2000. Pollinating efficacy of *Osmia cornuta* and *Apis mellifera* (Hymenoptera: Megachilidae, Apidae) on 'Red Delicious' apple. Environmental Entomology 29:235-240.

Vokou, D., T. Petanidou, and D. Bellos. 1990. Pollination ecology and reproductive potential of *Jankaea heldreichii* (Gesneriaceae); a Tertiary relict on Mt. Olympus, Greece. Biological Conservation 52:125-133.

Zaitoun, S., A. A. Al-Ghzawi, A. A. Al-Ghazawi, and A. Al-Qudah. 2007. Bee pollination and fruit set of Sumac (*Rhus coriaria*, Anacardiaceae) as a native herbal plant grown under semiarid Mediterranean conditions in Jordan. Advances in Horticultural Science:183-187.
